# Rod-shaped microglia interact with neuronal dendrites to regulate cortical excitability in TDP-43 related neurodegeneration

**DOI:** 10.1101/2024.06.30.601396

**Authors:** Manling Xie, Alessandra S. Miller, Praveen N. Pallegar, Anthony Umpierre, Yue Liang, Na Wang, Shuwen Zhang, Nagaswaroop Kengunte Nagaraj, Zachary C. Fogarty, Nikhil B. Ghayal, Björn Oskarsson, Shunyi Zhao, Jiaying Zheng, Fangfang Qi, Aivi Nguyen, Dennis W. Dickson, Long-Jun Wu

**Affiliations:** Department of Neurology, Mayo Clinic, Rochester, MN, USA; Center for Neuroimmunology and Glial Biology, Institute of Molecular Medicine, University of Texas Health Science Center, Houston, TX, USA; Department of Quantitative Health Sciences, Mayo Clinic, Rochester, MN, USA; Department of Neuroscience, Mayo Clinic, Jacksonville, FL, USA; Department of Laboratory Medicine and Pathology, Mayo Clinic, Rochester, Minnesota, USA

## Abstract

Motor cortical hyperexcitability is well-documented in the presymptomatic stage of amyotrophic lateral sclerosis (ALS). However, the mechanisms underlying this early dysregulation are not fully understood. Microglia, as the principal immune cells of the central nervous system, have emerged as important players in sensing and regulating neuronal activity. Here we investigated the role of microglia in the motor cortical circuits in a mouse model of TDP-43 neurodegeneration (rNLS8). Utilizing multichannel probe recording and longitudinal *in vivo* calcium imaging in awake mice, we observed neuronal hyperactivity at the initial stage of disease progression. Spatial and single-cell RNA sequencing revealed that microglia are the primary responders to motor cortical hyperactivity. We further identified a unique subpopulation of microglia, rod-shaped microglia, which are characterized by a distinct morphology and transcriptional profile. Notably, rod-shaped microglia predominantly interact with neuronal dendrites and excitatory synaptic inputs to attenuate motor cortical hyperactivity. The elimination of rod-shaped microglia through TREM2 deficiency increased neuronal hyperactivity, exacerbated motor deficits, and further decreased survival rates of rNLS8 mice. Together, our results suggest that rod-shaped microglia play a neuroprotective role by attenuating cortical hyperexcitability in the mouse model of TDP-43 related neurodegeneration.

## Introduction

Amyotrophic lateral sclerosis (ALS) is a progressive neurodegenerative disease that defined by its effect on motor neurons in brain and spinal cord, leading to progressive muscle weakness, paralysis, and eventually death. The cortical circuit, crucial for motor control, is intricately involved in ALS pathology ^1^. Clinical observations highlight cortical hyperexcitability as an early characteristic in ALS patients, which could contribute to early neuronal toxicity and motor neuron degeneration ^2^. Cortical hyperexcitability has also been observed in ALS mouse models ^3^ and other neurodegenerative diseases such as AD ^4,5^ and frontotemporal dementia ^6^. Identifying the intrinsic mechanisms that sense early aberrations within the motor cortical circuit may reveal novel therapeutic targets capable of rescuing motor neuron dysfunction and arresting disease progression in its early stages.

Microglia, as the principal immune cells surveilling the CNS microenvironment, play a critical role in maintaining brain homeostasis ^7,8^. Emerging evidence has highlighted the significant involvement of microglia in monitoring and regulating neuronal activity under both physiological and pathological conditions ^9,10^. Notably, both neuronal hyperactivity (*e.g.,* seizures) or hypoactivity (*e.g.,* anesthesia) increase microglial process dynamics and interactions with neurons ^11-13^. While microglial activation is acknowledged as a hallmark in ALS ^14^, their specific role in regulating motor cortical excitability remains largely unknown. Understanding the mechanism of microglia in sensing and modulating motor cortical excitability in ALS as well as their neuroprotective role could potentially lead to novel therapeutics.

In this study, we utilized multichannel probe recording coupled with chronic *in vivo* calcium imaging to monitor neuronal activity in the motor cortex of a mouse model of TDP-43 neurodegeneration, the rNLS8 mice. In this model, human TDP-43 without a nuclear localization sequence is expressed in a DOX-dependent manner ^15^. Upon initiating TDP-43 expression by removing the DOX diet, the mice exhibited characteristic motor deficits ^15,16^. This unique feature allowed us to precisely monitor neuronal activity during disease progression in real-time fashion. We observed neuronal hyperactivity and a unique microglia subpopulation, rod-shape microglia, in the cortex at the initial stages of disease progression in the rNLS8 mice. Rod-shaped microglia formed tight complexes with neuronal apical dendrites that may block excitatory synaptic inputs to attenuate cortical excitability. Our findings highlight the neuroprotective role of rod-shaped microglia in dampening neuronal excitability within the motor cortical circuits in TDP-43 neurodegeneration.

## Results

### Cortical hyperactivity occurs early in the motor cortex of rNLS8 mice

Hyperexcitability has been previously observed in the early stages of ALS progression in patients ^2^. To test whether this early hyperexcitable phenotype is recapitulated in the rNLS8 mice, we recorded single-unit, extracellular neuronal activity across cortical layers using a chronically implanted silicon probe (32-channels over 750 µm in primary motor cortex; **Fig. 1a**). We recorded neuronal activity first at the baseline (DOX ON: no TDP-43 overexpression) and then extended for a 4-week period of disease progression (DOX OFF: TDP-43 overexpression; **Fig. 1b**).

**Fig 1.**
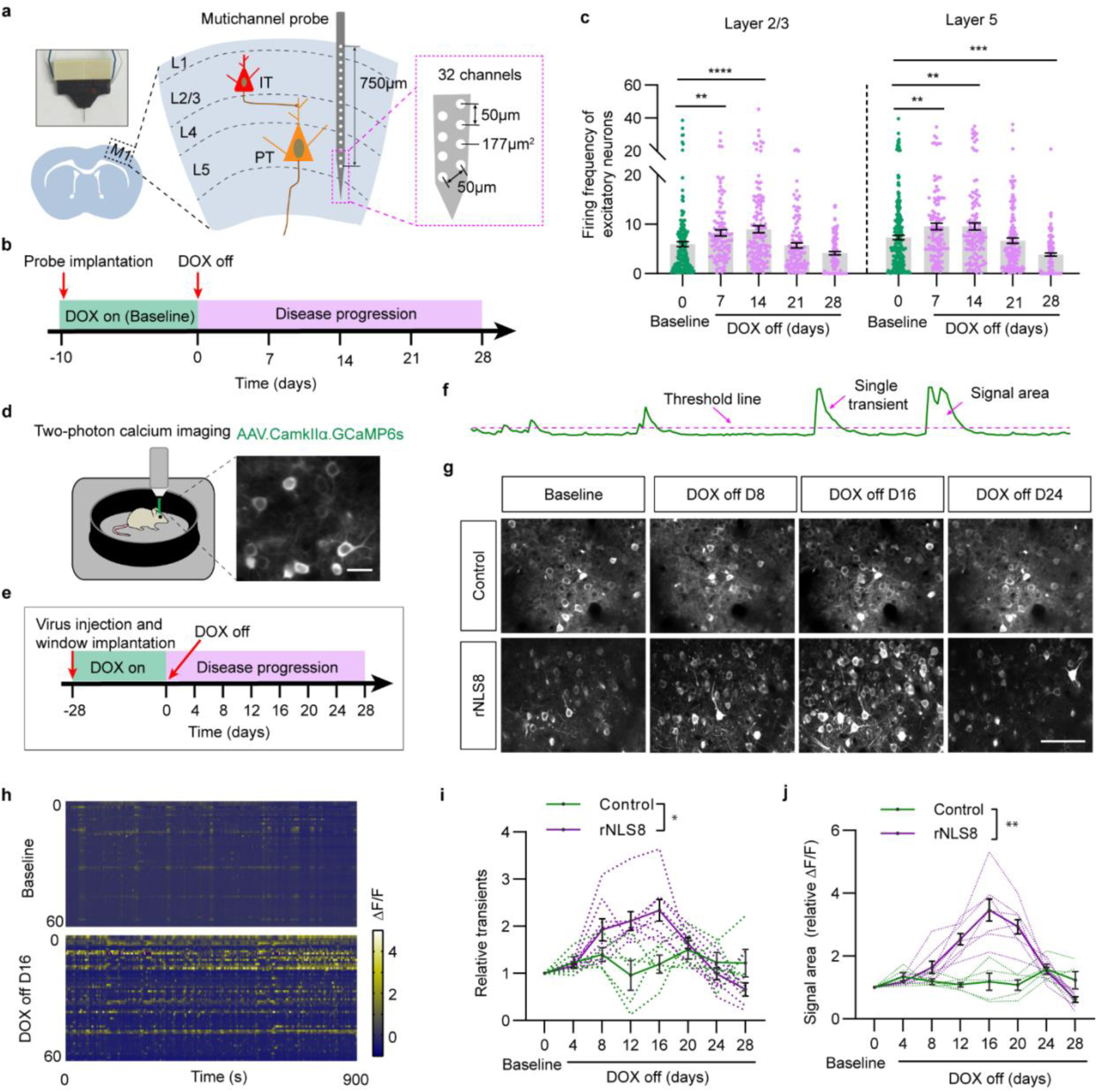
Motor cortical hyperactivity occurs early in the rNLS8 mice. **a**, Schematic illustration of in vivo recording using a silicon probe with 32 channels in the motor cortex of the control and rNLS8 mice. M1, Primary motor cortex; IT, intratelencephalic neuron; PT, pyramidal tract neuron. **b**, Experimental paradigm outlining probe implantation surgery and timeline for *in vivo* recording. After baseline recording, mice were switched to DOX-free chow to induce hTDP-43ΔNLS expression. **c**, Firing frequency of pyramidal neurons in layer 2/3 (179 neurons) and layer 5 (242 neurons) in the motor cortex of rNLS8 mice during baseline and disease progression (*n* = 7 mice). **d**, Schematic of *in vivo* two-photon Ca imaging in an awake, head-restrained mice with a representative average intensity projection image of neurons expressing CamkIIα-GCaMP6s. Scale bar, 20 µm. **e**, Experimental paradigm for AAV.CamkIIα.GCaMP6s virus injection and cranial window implantation surgery, with a timeline for chronic window imaging. **f**, Representative ΔF/F calcium traces from the soma of a neuron expressing CamkIIα-GCaMP6s, with the threshold line, single transient, and signal area of Ca activity indicated by pink arrows separately. **g**, Representative images of layer 2/3 neuronal calcium activity in control and rNLS8 mice during baseline and disease progression. Scale bars, 100 μm. **h**, Heatmap of calcium activity from 60 representative somatic ROIs in one animal under baseline and DOX off day 16. **i**, Quantification of transients (ratio) of neuronal calcium activity during baseline and disease progression (*n =* 5 per group). **j**, Quantification of ΔF/F·s signal area (ratio) of neuronal calcium activity during baseline and disease progression (*n =* 5 per group). Statistical analysis, one-way ANOVA followed by Fisher’s post hoc test (**c,** dot: one cell), two-way ANOVA followed by Sidak’s post hoc test (**i, j,** dotted lines represent longitudinal imaging regions. Error bars, mean ± s.e.m. NS = not significant; \**P* < 0.05; \*\**P* < 0.01; *** *P* < 0.001; **** *P* < 0.0001.

Distinctions between excitatory and inhibitory neurons were made based on spike duration combined with interspike interval (ISI) histograms (**Extended Data Fig. 1a-f and see Methods**). The probe of 32 channels successfully covered neuronal activities from all cortical layers (**Extended Data Fig. 1g**). At baseline, neuronal firing frequency was similar between genotypes in both layer 2/3 and layer 5 (**Extended Data Fig. 1h**). As expected, inhibitory neurons showed a higher firing frequency compared to excitatory neurons at baseline (**Extended Data Fig. 1h**). In addition, neuronal activity was stable in control mice over consecutive recoding days, signifying the reliability of the recording system (**Extended Data Fig. 1i,j**). In rNLS8 mice, we observed that the firing frequency of excitatory neurons increased during the first 2 weeks post-DOX diet removal in both layer 2/3 and layer 5 (**Fig. 1c**). To corroborate the findings from the silicon probe recordings, we employed longitudinal *in vivo* two-photon calcium imaging approaches (**Fig. 1d,e**). We transfected rNLS8 mice and their controls with AAV.CaMKIIa.GCaMP6s (**Fig. 1d,e**) and calcium activity of excitatory neurons was then imaged in the primary motor cortex (layer 2/3) through a chronic cranial window. Neuronal activity was estimated by analyzing the ΔF/F·s signal area of calcium transients with baseline normalization (**Fig. 1f and see Methods**). In alignment with silicon probe findings, neuronal calcium transient frequency gradually increased following DOX diet removal, reaching peak elevations 2-3 weeks post-DOX diet removal (**Fig. 1g-i and Supplementary Video 1,2**). Correspondingly, calcium signal area also showed a sustained increase before reduction (**Fig. 1j**). At their peak, these neurons exhibited a 228 ± 49% increase in the average ΔF/F·s calcium signal area (**Fig. 1j**). Control mice exhibited consistent calcium activity of excitatory neurons over the 4-week recoding period (**Fig. 1g-j, and Supplementary Video 3,4**). These observations indicate that rNLS8 mice experience cortical hyperexcitability during the early stage of TDP-43 overexpression.

We next explored the potential mechanisms underlying hyperactivity in pyramidal neurons. Specifically, we examined the level of neurodegeneration, the electrophysiological properties of pyramidal neurons, and the contribution of inhibitory neurons. NeuN staining revealed no significant changes in neuron density up to 3 weeks post-DOX diet removal **(Extended Data Fig. 2a,b**). However, Nissl staining showed a significant reduction in Nissl bodies within pyramidal neurons as the disease progressed, indicating neuronal stress (**Extended Data Fig. 2c,d**). Next, we conducted electrophysiological recordings of pyramidal neurons 16 days post-DOX diet removal, a time point coinciding with the peak of neuronal hyperactivity. These recordings indicated that pyramidal neurons in rNLS8 mice exhibited increased excitability compared to control mice (**Extended Data Fig. 2e-g**). Silicon probe recordings indicated that the firing frequency of inhibitory neurons remained largely unchanged in the first two weeks, suggesting that changes in inhibitory neurons are not the direct trigger for early-stage cortical hyperactivity (**Extended Data Fig. 2h,i**). Altogether, these data demonstrate cortical hyperactivity within the motor cortex of rNLS8 mice at early stages of disease progression, which is associated with increased excitability of pyramidal neurons.

### Spatial transcriptomics analysis reveals cortical microglia as the major responder

To explore the broad picture of cellular changes in the brain of rNLS8 mice in response to neuronal hyperactivity, we performed spatial RNA sequencing (spRNA-seq, 10x Visium) to create transcriptomic maps across various brain regions. Specifically, we collected two coronal sections from both rNLS8 and control mice three weeks post-DOX diet removal, a time point selected to capture peak neuronal hyperactivity (**Fig. 2a**). Each section provided over 1,700 transcriptomic profiles from individual spots, totaling 8,474 profiles with a median depth of 8,619 unique molecular identifiers (UMIs) per spot and an average of 3,754 genes per spot (**Extended Data Fig. 3a-e**). By aligning spatial gene expressions with immunofluorescence staining (NeuN/DAPI), we confirmed spatial concordance for the transcriptomic data (**Fig. 2b**). Using a clustering-based approach, we annotated the regional identity of Visium spots, classifying them into 6 major regions, including the cerebral cortex (CTX, layers I to V), cortical subplate (CTXsub), olfactory areas (OLF), white matter (WM), striatum (STR) and pallidum (PAL), based on the Allen Mouse Brain Atlas ^17^ (**Fig. 2c-e, Extended Data Fig. 3f, Supplementary file 1**). Notably, we identified a unique cortical cluster in the rNLS8 mice, termed the ‘sp-rNLS8-unique cluster’ (**Fig. 2c-e**). All the clusters were consistently captured across each sample (**Extended Data Fig. 4a-d**).

**Fig 2.**
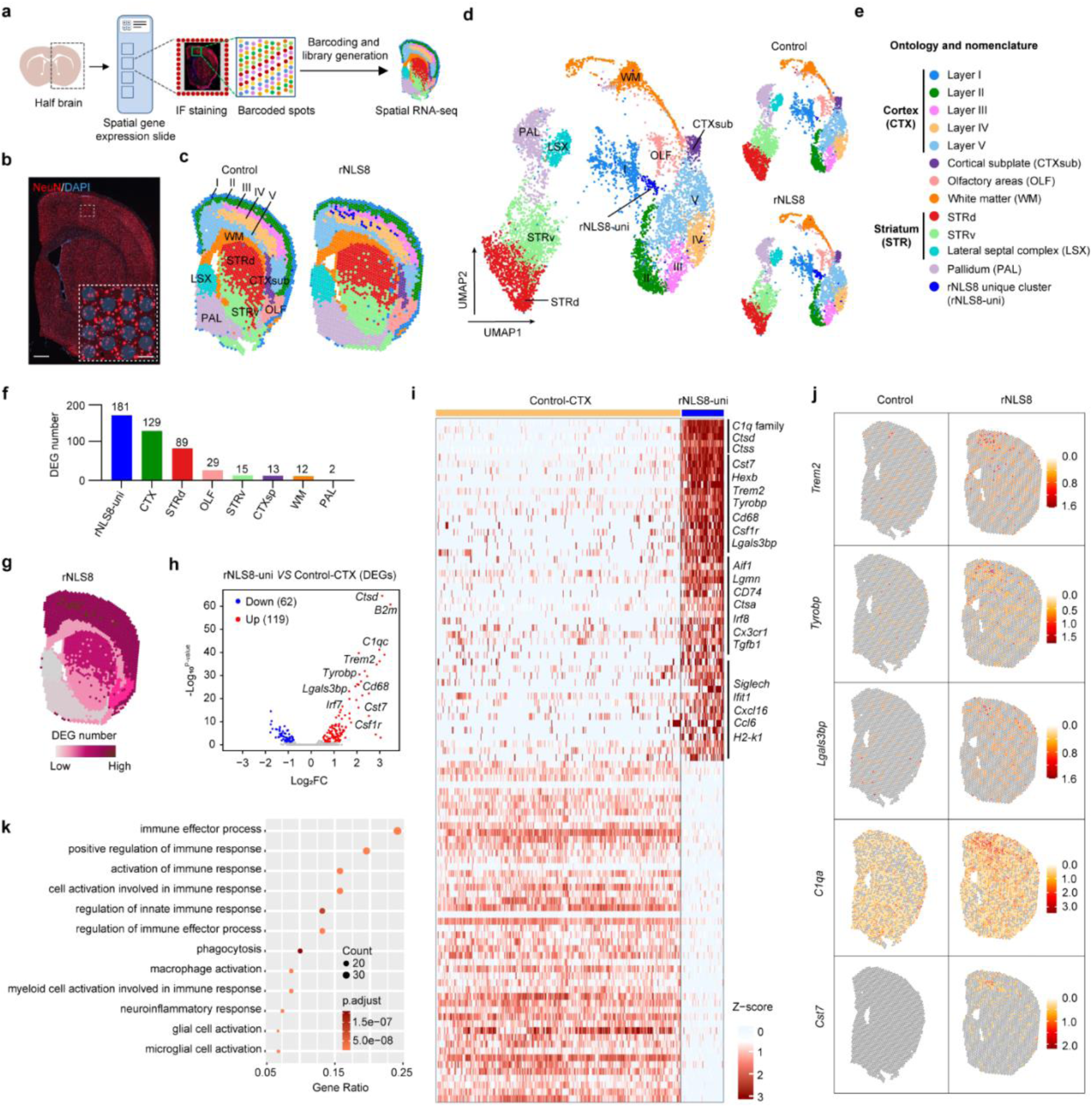
Spatial transcriptomics analysis of the cortical area reveals microglia as the major responder. **a**, Schematic workflow for spatial RNA sequencing (spRNA-seq). Mice were euthanized on 3 weeks post-DOX diet removal. Two coronal brain tissue sections from control or rNLS8 mice were processed for 10X Visium spatial transcriptomics. **b,** Immunostaining of NeuN and DAPI in the 10X Visium slices. Scale bar, 300 µm. Inset shows NeuN/DAPI expression with the barcoded spots at higher magnification as indicated by the area in dotted white box in the motor cortex. Scale bar, 100 µm. **c, d,** Representative spatial transcriptome data for a control and rNLS8 mouse, colored by cluster-level annotation and represented as spatial transcriptome (**c**) and UMAP (**d**). Cluster-level data determined by Seurat clustering. **e,** Data description and abbreviations of ontology and nomenclature for spatial transcriptome data. **f,** Differential gene expression analysis comparing control and rNLS8 mice across all brain regions delineated with spatial transcriptomics. **g,** Visualization of the number of differentially expressed genes (DEGs) in the corresponding brain regions. **h,** Volcano plot of DEGs in ‘rNLS8 unique cluster’ from spRNA-seq data. The cortex (CTX) cluster (combining all cortical layers) in the control group was served as baseline. Significance is indicated at adjusted *P* value ≤ 0.05 and |avg_log2FC| ≥ 0.5 (log2FC, log2-fold change). **i,** Heatmap of DEGs in ‘rNLS8 unique cluster’ from spRNA-seq data. **j,** Spatially resolved expression of select genes across tissue sections from control and rNLS8 mice. **k,** Gene Ontology (GO) enrichment analysis was performed for the ‘rNLS8 unique cluster’ from spRNA-seq data. Dot plot of the selected GO terms in order of gene ratio. The size of each dot indicates the number of genes in the significant DEG list that are associated with each GO term, and the color of the dots corresponds to the adjusted *P*-values.

We then analyzed differentially expressed genes (DEGs) to explore the transcriptional changes across all identified regions between the rNLS8 group and the control group. The CTX cluster (combining all cortical layers) in the control group served as baseline to identify DEGs in the ‘sp-rNLS8-unique cluster’. We identified a total of 470 DEGs across all clusters, with the ‘sp-rNLS8-unique cluster’ and CTX cluster showing more pronounced regulation of DEGs compared to other regions (**Fig. 2f,g and Supplementary file 2**), highlighting the cortex as the most significantly affected area in the brain. Despite its small size, the ‘sp-rNLS8-unique cluster’ exhibited the most significant number of DEGs (**Fig. 2f-h and Supplementary file 2**). Notably, the majority of the top 100 upregulated DEGs in this cluster were highly or exclusively expressed in microglia based on the brain RNAseq database (https://brainrnaseq.org), including members of the complement *C1q* family, *Cst7*, *Hexb*, *Trem2*, *Tyrobp*, *Cd68*, *Csf1r*, *Lgals3bp*, *H2-k1*, *Alx*, and *Ifitm3*, among others (**Fig. 2i and Supplementary file 2**). These microglia-specific genes were further evaluated anatomically through spRNA-seq (**Fig. 2j**). Additionally, Gene Ontology (GO) enrichment analysis confirmed that ‘microglia activation’ was one of the most upregulated functional terms, along with other terms related to ‘immune responses’, ‘phagocytosis’ and ‘neuroinflammatory response’ (**Fig. 2k**). DEGs and GO enrichment analysis from the CTX cluster were consistent with those from ‘sp-rNLS8-unique cluster’, strongly indicating microglia/macrophage recruitment or activation (**Extended Data Fig. 4e-g**). Additionally, we observed a subset of differentially expressed neuronal-related genes highly upregulated in the CTX cluster, predominantly enriched in GO functional terms such as ‘synaptic transmission’, ‘synapse organization’ and ‘synapse pruning’ (**Extended Data Fig. 4e-g**). These findings align with our electrophysiology data of increased neuronal excitability in rNLS8 mice. Overall, our spatial RNA-seq data reveal significant transcriptional reprogramming in the cortical regions in response to neuronal hyperactivity, with microglia emerging as primary responders.

### Microglia subpopulation identified in rNLS8 mice and ALS patients: rod-shaped microglia

To further assess the transcriptional profile changes induced by neuronal hyperactivity in cortical regions, we conducted single cell RNA sequencing (scRNA-seq) (**Fig. 3a**). We obtained high-quality single-cell gene expression profiles (54,607 from 4 rNLS8 mice and 46,555 from 4 control mice), detecting a median of 6483 (rNLS8), 5901(control) unique mRNA molecules and 2163 (rNLS8), 2081(control) genes per cell. Through unsupervised clustering, we identified 13 distinct clusters based on well-defined marker genes, representing major cell types (**Fig. 3a,b)**. These include microglia, vascular endothelium, astrocytes, neurons, ependymal cells, endothelial cells, choroid plexus cells, smooth muscle cells, pericytes, monocytes/macrophages, mitotic microglia, lymphocytes, granulocytes, and fibroblasts (**Fig. 3a-c, Extended Fig. 5a and Supplementary file 3**). The clustering was not biased by individual samples or genders (**Extended Fig. 6a-c**). UMAPs of all cell types from the scRNA-seq data again corroborated microglial transcriptional reprogramming in rNLS8 mice compared with control mice (**Fig. 3d**). Additionally, a significant presence of the mitotic microglia cluster was observed in the rNLS8 group (**Fig. 3b,d and Extended Fig. 6d,e**), indicating a highly proliferative microglia population at this stage of disease progression.

**Fig 3.**
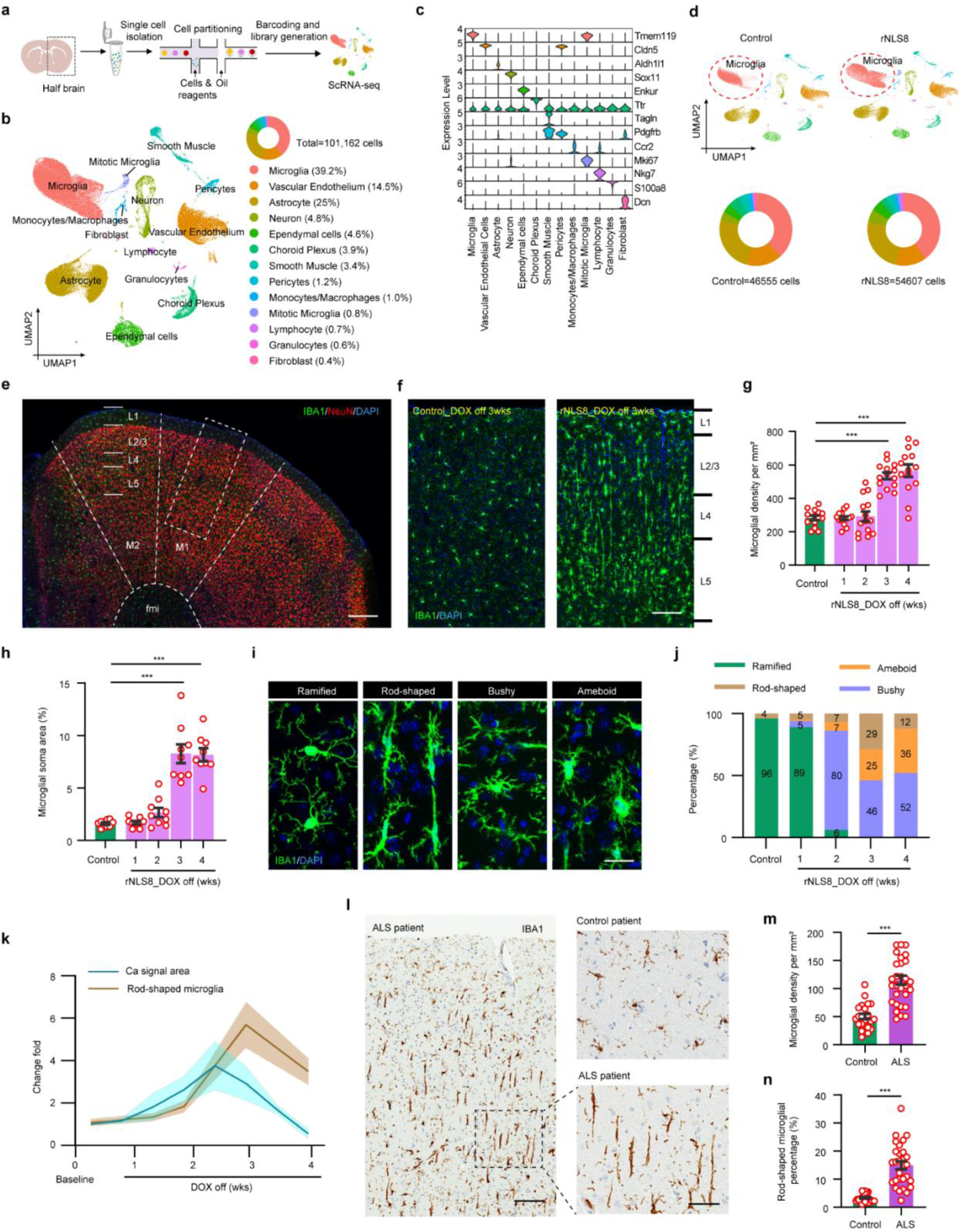
A unique microglia subpopulation identified in rNLS8 mice: rod-shaped microglia. **a**, Schematic workflow for single cell RNA sequencing (scRNA-seq). Mice were euthanized at 3 weeks post-DOX diet removal. Four half brains from control or rNLS8 mice were processed for scRNA-seq. **b,** UMAP visualization of all cells identified by scRNAseq. Cells are color-coded by their identities (number of cells = 101,162 from 8 samples). Pie charts show the cell proportions of each cluster. **c,** Stacked violin plot of known marker genes for each identified cell type. **d,** UMAP visualization of all cells identified by scRNA-seq in control and rNLS8 mice. Pie charts show the cell proportions of each cluster. **e**, Layer structure of the motor cortex of a control mouse brain shown by NeuN and IBA1 expression. The primary motor cortex (M1), supplementary motor area (M2), and lateral ventricle (LV) are separated by a dashed line. Scale bar: 300 µm. **f,** IBA1 expression in M1 of control and rNLS8 mice at 3 weeks post-DOX diet removal at higher magnification, as indicated by the area in dotted white box in **e**. Scale bar, 100 µm. **g, h,** Quantification of microglial density (**g,** *n* = 14 per group) and soma area (**h,** *n* = 9 per group) in M1 of control and rNLS8 mice at 3 weeks post-DOX diet removal. **i**, Representative images of microglia with different morphologies, including ramified microglia identified at baseline, and rod-shaped microglia, bushy microglia, and amoeboid microglia identified at 3 weeks post-DOX diet removal. Scale bar, 10 µm. Bushy microglia are characterized by an enlarged soma (1 to 2 times the diameter of the soma in ramified cells) with short and poorly ramified processes. Ameboid microglia have a larger cell body (more than twice the diameter of the soma in ramified cells) with one or without any process. Rod-shaped microglia are characterized by a slender, elongated cell body and few planar processes, which are distinctly polarized. **j**, Percentage of microglia with different morphologies during baseline and disease progression. **k,** Correlation of rod-shaped microglia percentage change fold with cortical neuronal Ca signal area change fold. **l**, Representative images of IBA1 staining in motor cortex tissue of ALS patients (*n* = 31) and age-matched controls (*n* = 25). Scale bar, 100 µm. High-magnification images as indicated by the areas in the dotted black boxes are shown on the right. scale bar, 50 µm. **m, n,** Quantification of microglia density (**m**) and rod-shaped microglia percentage (**n**) in the motor cortex of ALS patients and age matched controls. Statistical analysis, one-way ANOVA followed by Fisher’s post hoc test (**g, h**), or two-tailed unpaired Student’s t-test (**m, n**). Error bars, mean ± s.e.m. NS = not significant, \**P* < 0.05; \*\**P* < 0.01; *** *P* < 0.001; **** *P* < 0.0001.

To validate the observations from both spRNA-seq and scRNA-seq, we analyzed changes in microglia using IBA1 staining. In control mice, microglia were uniformly distributed throughout the motor cortical layers (**Fig. 3e,f**). However, in rNLS8 mice, we observed a significant increase in microglial density and soma size within the motor cortex as the disease progressed, with a peak at three weeks post-DOX diet removal (**Fig. 3f-h**). Unlike the ramified microglia seen in a homeostatic state, activated microglia exhibited a reactive phenotype with diverse morphologies (**Fig. 3i**), including amoeboid microglia, bushy microglia, and distinct rod-shaped microglia.

Rod-shaped microglia are characterized by a slender, elongated cell body and few planar processes, which are distinctly polarized (**Fig. 3i**). They were mainly distributed in layer 2/3 and layer 4 (**Fig. 3f**). Notably, quantitative analysis revealed that the percentage of rod-shaped microglia peaked three weeks post-DOX diet removal, coinciding with the peak of neuronal hyperactivity (**Fig. 3j,k**). This suggests that rod-shaped microglia are the primary responders to neuronal hyperactivity. To validate the existence of rod-shaped microglia in ALS patients, we performed immunohistochemistry staining of microglia in the motor cortex of postmortem tissues. We found rod-shaped microglia were abundant in ALS patients compared to controls, as well as the overall microglial density (**Fig. 3l-n and Supplementary file 4**). Altogether, these findings highlight that a unique subpopulation of microglia—rod-shaped microglia—responds to cortical hyperactivity in a temporally and spatially specific manner.

### Rod-shaped microglia exhibit unique transcriptional and functional profiles

To explore the transcriptomic signature of microglia, particularly rod-shaped microglia, in rNLS8 mice, we re-clustered microglia from combined clusters across all groups, identifying 7 distinct microglia subpopulations from a total of 39,645 cells (**Fig. 4a**). The clustering was unbiased by individual samples or genders (**Extended Fig.7a-d**). These subpopulations were characterized using lineage-specific markers (**Extended Fig.7e and Supplementary file 5**). Clusters 0, 1, and 3 (MG0, MG1, and MG3), predominantly found in the control group, were identified as ‘homeostatic microglia’, based on their higher expression of homeostatic marker genes such as *P2ry12*, *Tmem119*, *Cx3cr1*, and *Hexb* (**Fig. 4b,c, Extended Fig.7e and Supplementary file 5**). Cluster MG6, with a slight but non-significant increase in rNLS8 mice, was enriched with genes related to cell cycle, including members of the *Mcm* gene family, *Dtl*, and *Smc1b* (**Fig. 4b,c, Extended Fig.7e and Supplementary file 5**). Clusters 2, 4, and 5 (MG2, MG4, and MG5) were significantly enriched in the rNLS8 group and classified as ‘sc-rNLS8-specific microglia clusters’ (**Fig. 4b,c**). Notably, MG2 exhibited similarities with ‘disease-associated microglia’ (DAM) identified in AD mouse models ^18,19^, characterized by the downregulation of homeostatic genes and upregulation of DAM marker genes (such as *Apoe*, *Alx*, *Lpl*, *Spp1*, *Igf1*), as well as the *Lgals* family genes and cytokine/chemokine genes, and was thus classified as ‘sc-rNLS8-DAM’ (**Fig. 4b,c, Extended Fig.7e,f and Supplementary file 5**). MG4 resembled homeostatic clusters and exhibited very limited marker genes, representing a transitional state (**Extended Fig.7e and Supplementary file 5**). MG5, identified as ‘interferon-responsive microglia’, showed high expression of *Ifit* family genes (**Fig. 4b,c, Extended Fig.7e,f and Supplementary file 5**).

**Fig 4.**
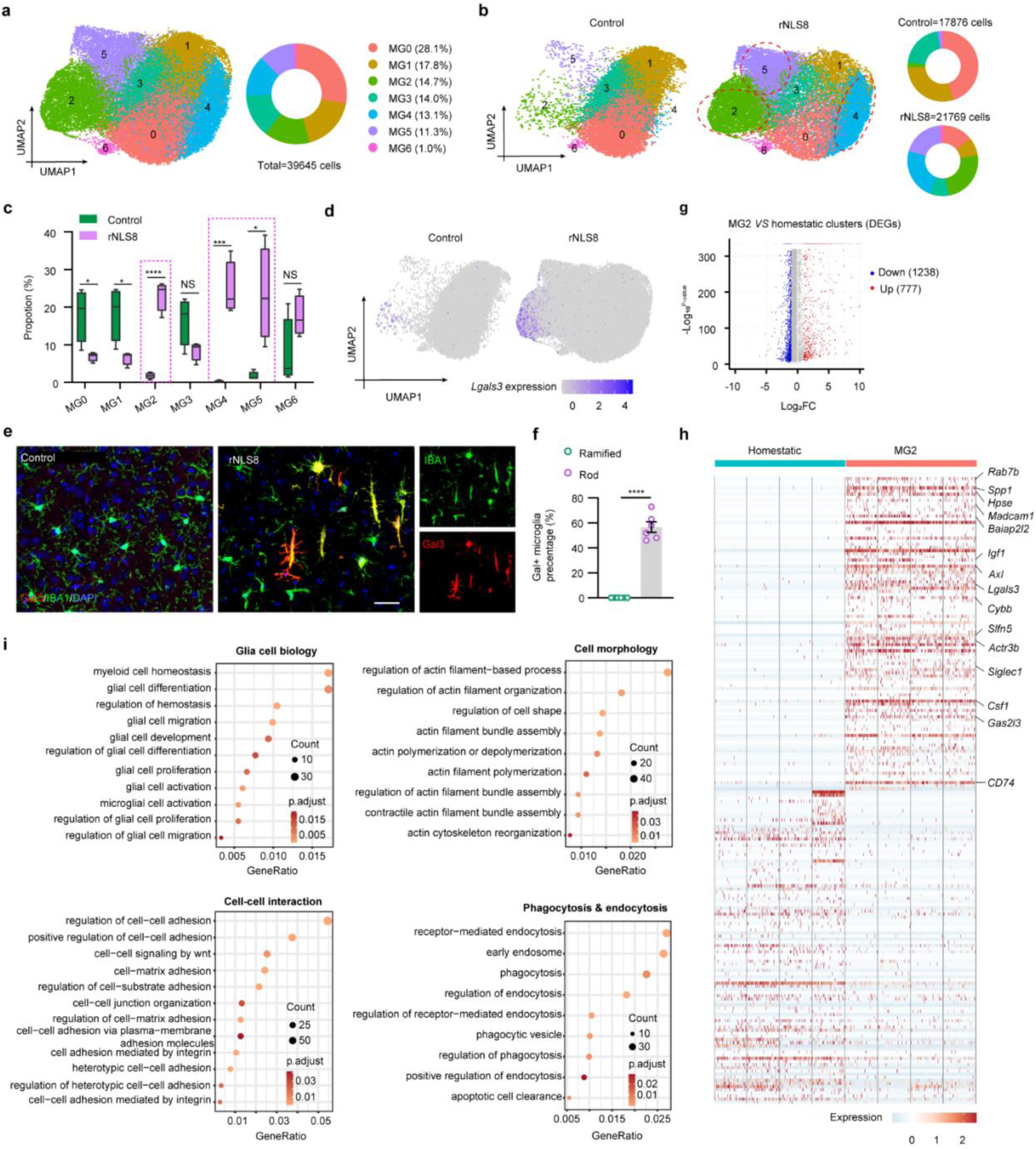
Rod-shaped microglia exhibit a unique transcriptional and functional profile. **a,** UMAP visualization of microglia subclusters identified by scRNA-seq. Cells are color-coded by their identities (number of cells = 39,645 from 8 samples). Pie charts show the cell proportions of each cluster. **b,** UMAP visualization of microglia subclusters in control and rNLS8 mice. Pie charts show the cell proportions of each cluster. **c,** Quantification of cell proportions of each microglial subcluster in control and rNLS8 mice. **d,** Feature plot of *Lgals3* expression in control and rNLS8 mice from scRNA-seq analysis. **e, f,** Representative images (**e**) and quantification (**f**) of Galactin 3 (Gal3, red) positive microglia (IBA1, green) in M1 of control and rNLS8 mice at 3 weeks post-DOX diet removal. scale bar, 50 µm. **g,** Volcano plot of DEGs in microglia subcluster 2 (MG2). Homeostatic microglia cluster (combination of MG0, MG1 and MG3 from the control group) served as baseline. Significance at adjusted *P* value ≤ 0.05 and |avg_log2FC| ≥ 1 (log2FC, log2-fold change). **h,** Heatmap of top 100 upregulated and downregulated DEGs in the MG2 cluster compared with the homeostatic microglia cluster. **i,** Dot plot of the selected GO terms in order of gene ratio in MG2. The size of each dot indicates the number of genes in the significant DEG list that are associated with each GO term, and the color of the dots corresponds to the adjusted P-values. Statistical analysis, two-tailed unpaired Student’s t-test (**c**). Error bars, mean ± s.e.m. NS = not significant, \**P* < 0.05; \*\**P* < 0.01; *** *P* < 0.001; **** *P* < 0.0001.

To determine which cluster rod-shaped microglia belonged to, we screened marker genes from ‘sc-rNLS8-specific microglia clusters’ using immunostaining. Our results showed that Galectin-3 (encoded by *Lgals3*) specifically labeled approximately 60% of rod-shaped microglia, but not other markers, like MHC II, LPL, CD11c or AXL (**Fig. 4d-f and Extended Fig. 8a-d**). This indicates that rod-shaped microglia are part of the ‘sc-rNLS8-DAM cluster’ (MG2). From this insight, we conducted a DEG analysis for MG2 to explore the potential functions of rod-shaped microglia. The homeostatic clusters (MG0, MG1, and MG3 combined) in the control group served as a control. A total of 2,015 significant DEGs were identified in MG2 (**Fig. 4g, and Supplementary file 6**). Of note, analysis of the top 100 upregulated DEGs revealed upregulation of genes involved in cell adhesion (*Madcam1*, *Lgals3*, *Siglec1*), phagocytosis (*Spp1*, *Axl*, *Cybb*), and cell remodeling (*Baiap2l2*, *Actr3b*, *Gas2l3*) (**Fig. 4h**). GO enrichment analysis further revealed that the top enriched processes related to ‘glial cell activation’, ‘regulation of cell morphology’, ‘cell-cell adhesion’, and ‘phagocytosis & endocytosis’ (**Fig. 4i**), suggesting intriguing undiscovered function of this microglia cluster. Thus, microglia subcluster analysis from scRNA-seq extends spRNA-seq by revealing the unique transcriptional and functional profiles of rod-shaped microglia in rNLS8 mice.

### Rod-shaped microglia interact with neuronal dendrites

To evaluate the function of rod-shaped microglia, we hypothesized that this subpopulation might interact with key brain structures in the cortex. This hypothesis is supported by the highlighted GO terms from MG2, the distinctive morphology of the rod-shaped microglia, and their specific distribution across cortical layers. Co-immunostaining of IBA1 with markers for blood vessels (CD31), axons (ANKG), and dendrites (MAP2) showed that about 70% of interactions by rod-shaped microglia were with dendrites, rather than with blood vessels or axons (**Fig. 5a-f**). Additionally, we crossed rNLS8 mice with Thy1-YFP mice to specifically label layer 5 pyramidal neurons and found that nearly 50% of these rod-shaped microglia are engaged with apical dendrites (**Fig. 5g-j**). This contrasted with only about 10% interaction observed in the ramified microglial soma of control mice (**Fig. 5g-j**). Ultrastructural studies in rNLS8 mice identified three distinct interaction patterns between rod-shaped microglia and neuronal dendrites: Type 1, where the microglia processes connect to both apical dendrites and spines; Type 2, involving microglial body contacting apical dendrites; and Type 3, where microglia completely encase apical dendrites (**Fig. 5h-i and Supplementary Video 5**).

**Fig 5.**
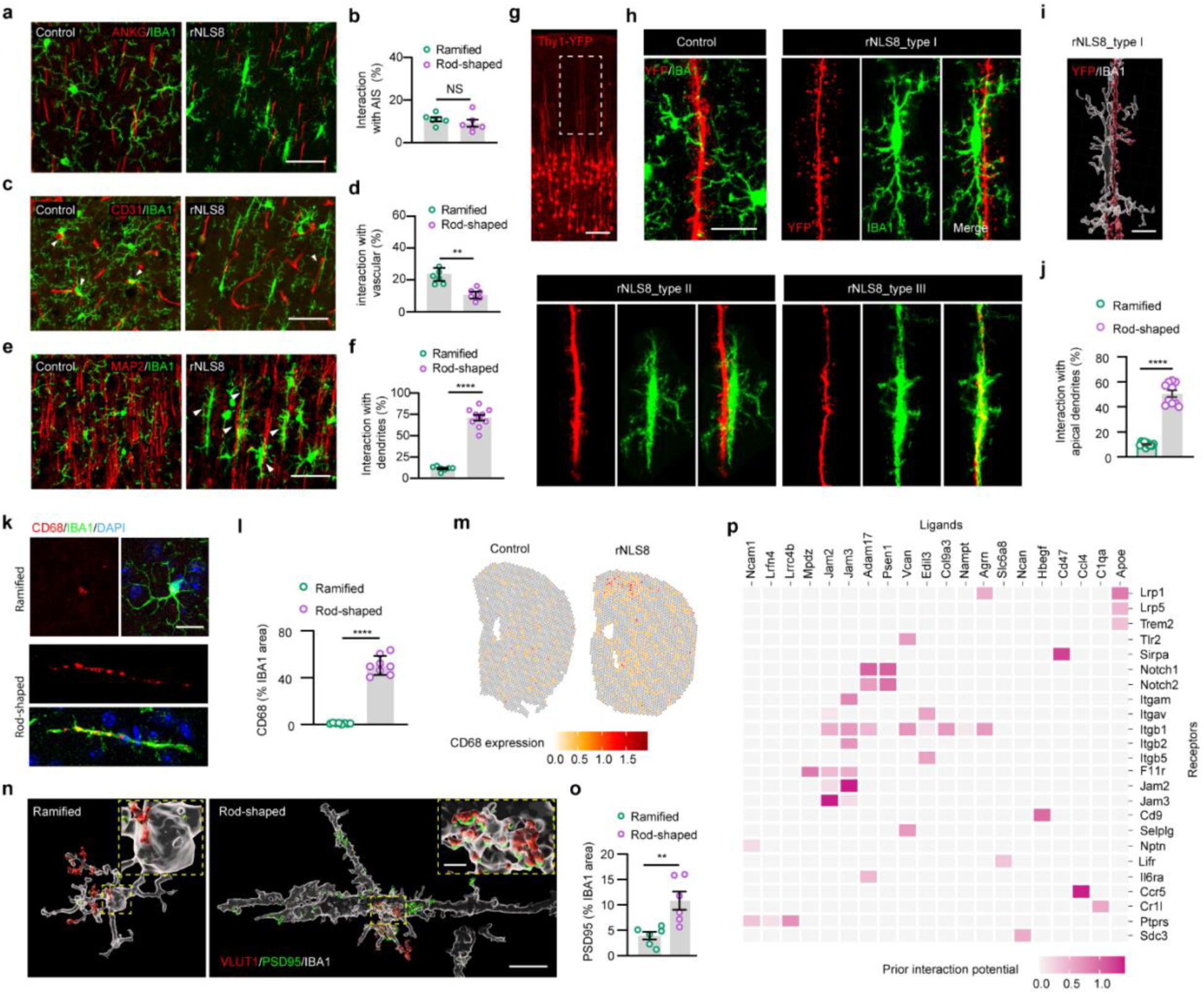
Rod-shaped microglia interact with neuronal dendrites. **a,b,** Representative images of co-staining of microglia (IBA1, green) with axon initial segment (AIS) (ANKG, red) (**a**) and quantification (**b**) of their interaction in M1 of control and rNLS8 mice at 3 weeks post-DOX diet removal. Scale bar, 50 µm. **c,d,** Representative images of co-staining of microglia (IBA1, green) with blood vessel (CD31, red) and quantification of their interaction (**c**) in M1 of control and rNLS8 mice at 3 weeks post-DOX diet removal. The interactions are indicated by arrowheads. Scale bar, 50 µm. **e,f,** Representative images of co-staining of microglia (IBA1, green) with neuronal dendrites (MAP2, red) (**e**) and quantification (**f**) of their interaction in M1 of indicated groups at 3 weeks post-DOX diet removal. The interactions are indicated by arrowheads. Scale bar, 50 µm. **g**, Representative images of YFP positive pyramidal neurons in M1 of thy1-YFP mice. Apical dendrites are indicated by the area in dotted white box. Scale bar, 100 µm. **h,** Representative images of microglia (IBA1, green) interaction with apical dendrites (red) in M1 of control and rNLS8:Thy1-YFP mice at 3 weeks post-DOX diet removal. Scale bar, 20 µm. **i,** Representative 3D-image of microglial (purple) interaction with apical dendrites (green). Scale bar, 10 μm. **j**, Quantification of ramified microglia (control) and rod-shaped microglia (rNLS8) interaction with apical dendrites in M1 at 3 weeks post-DOX diet removal (*n* = 10 per group). **k,l,** Representative images (**k**) and quantification (**l,** *n* = 8 per group) of CD68 (red) expression in ramified (control) and rod-shaped microglia (rNLS8) at 3 weeks post-DOX diet removal. Scale bar, 20 µm. **m,** Spatially resolved expression of *Cd68* across tissue sections from control and rNLS8 mice at 3 weeks post-DOX diet removal. **n,** Representative 3D-image of microglial (IBA1, white) phagocytosis of excitatory synaptic materials (VLUT1/PSD95, red/green). Scale bar, 20 μm. Insets show images at higher magnification as indicated by the area in dotted yellow boxes. Scale bar, 5 µm. **o**, Quantification of ramified (control) and rod-shaped microglial (rNLS8) phagocytosis of PSD95 in M1 at 3 weeks post-DOX diet removal (*n* = 6 per group). **p**, NicheNet’s ligand–target matrix denotes the regulatory potential between neuron-ligands and target receptors from microglia cluster. Statistical analysis, two-tailed unpaired Student’s t-test (**b, d, f, j, l and o**). Error bars, mean ± s.e.m. NS = not significant, \**P* < 0.05; \*\**P* < 0.01; *** *P* < 0.001; **** *P* < 0.0001.

We next explored whether the proximity of rod-shaped microglia to dendrites influences excitatory synaptic inputs. Consistent with the highlighted GO terms, upregulation of CD68 and downregulation of homeostatic markers (P2Y12 and TMEM119) confirmed the reactive, phagocytic nature of rod-shaped microglia (**Fig. 5k-m and Extended Data Fig. 9a,b**). Critically, we found that rod-shaped microglia extensively engulfed excitatory synapses, suggesting they might modulate neuronal activity by eliminating these synaptic inputs (**Fig. 5n,o and Supplementary Videos 6 and 7**). To better understand the mechanisms mediating the interactions between rod-shaped microglia and neurons, we conducted ligand-receptor interactome analyses of microglia and neuron clusters from scRNA-seq data using NicheNet ^20^. The results showed that the highly expressed receptors on microglia were specifically associated with cell adhesion (*Notch* family genes, *Integrin* family genes, *Nptn*, *Jam2/3*, *F11r*) and phagocytosis (*Trem2*, *Tlr2*, *Lrp1/5*) (**Fig. 5p and Extended Data Fig. 9c-e**). *Apoe* from neurons is predicted to be a potential ligand for the phagocytosis receptor (**Fig. 5p and Extended Data Fig. 9c-e**). The potential ligands for the cell adhesion receptors included *Ncam1*, *Lrfn4*, *Mpdz* and others (**Fig. 5p and Extended Data Fig. 9c-e**). Altogether, our findings suggest that rod-shaped microglia have a close physical relationship to the dendritic compartment and could play a crucial role in regulating the activity of pyramidal neurons by synaptic remodeling.

### TREM2/DAP12 axis regulates the formation of rod-shaped microglia and neuronal hyperactivity

We next wanted to identify the potential regulators mediating the transformation of rod-shaped microglia. Specifically, we performed RNA velocity analysis to detect critical genes that regulate dynamic transcriptional transitions among microglial clusters, with a specific focus on MG2 and its adjacent counterparts. This technique uses the ratio of unspliced pre-mRNA to spliced mRNA to infer the dynamic state of gene expression, characterizing the direction and rate of changes in cellular transcriptional states and enabling the deduction of trajectories among neighboring clusters^21,22^.

Our results showed that RNA velocity vectors originated from homeostatic clusters (MG0, MG1, MG3, and MG4) and pointed toward MG5 and MG6, eventually converging on MG2 (**Fig. 6a**). This suggests a continuous cell state transition trajectory. A dramatically increased latent time and pseudo-time, combined with the highest proportion of spliced RNA (74%) observed in MG2, confirmed that this cluster represents a terminal differentiation state (**Fig. 6b,c and Extended Data Fig. 10a,b**). We further explored the critical genes driving these microglial transitions. The results showed that the most dynamically expressed genes included those encoding transcriptional factors (e.g., *Zfhx3*, *Atf3*) and regulating ribosome function (e.g., *Rps21*, *Rpl13a*, *Rps3*) (**Fig. 6d, Extended Data Fig. 10c,d and Supplementary file 7**). Interestingly, *Tyrobp*, encoding the DNAX-activating protein of 12 kDa (DAP12), was also among the highly dynamic genes (**Fig. 6d,e and Supplementary file 7**). As the major adapter protein for TREM2, DAP12 forms a complex with TREM2 and acts as the principal regulator mediating the conversion of homeostatic microglia into DAM ^16^. Both *Tyrobp* and *Trem2* exhibited significant upregulation at later stages of pseudo-time, highlighting the crucial role of this signaling pathway in the microglial transition process (**Fig. 6e,f**). Collectively, these observations suggest that the TREM2/DAP12 axis may play an important role in the transformation of rod-shaped microglia in rNLS8 mice.

**Fig 6.**
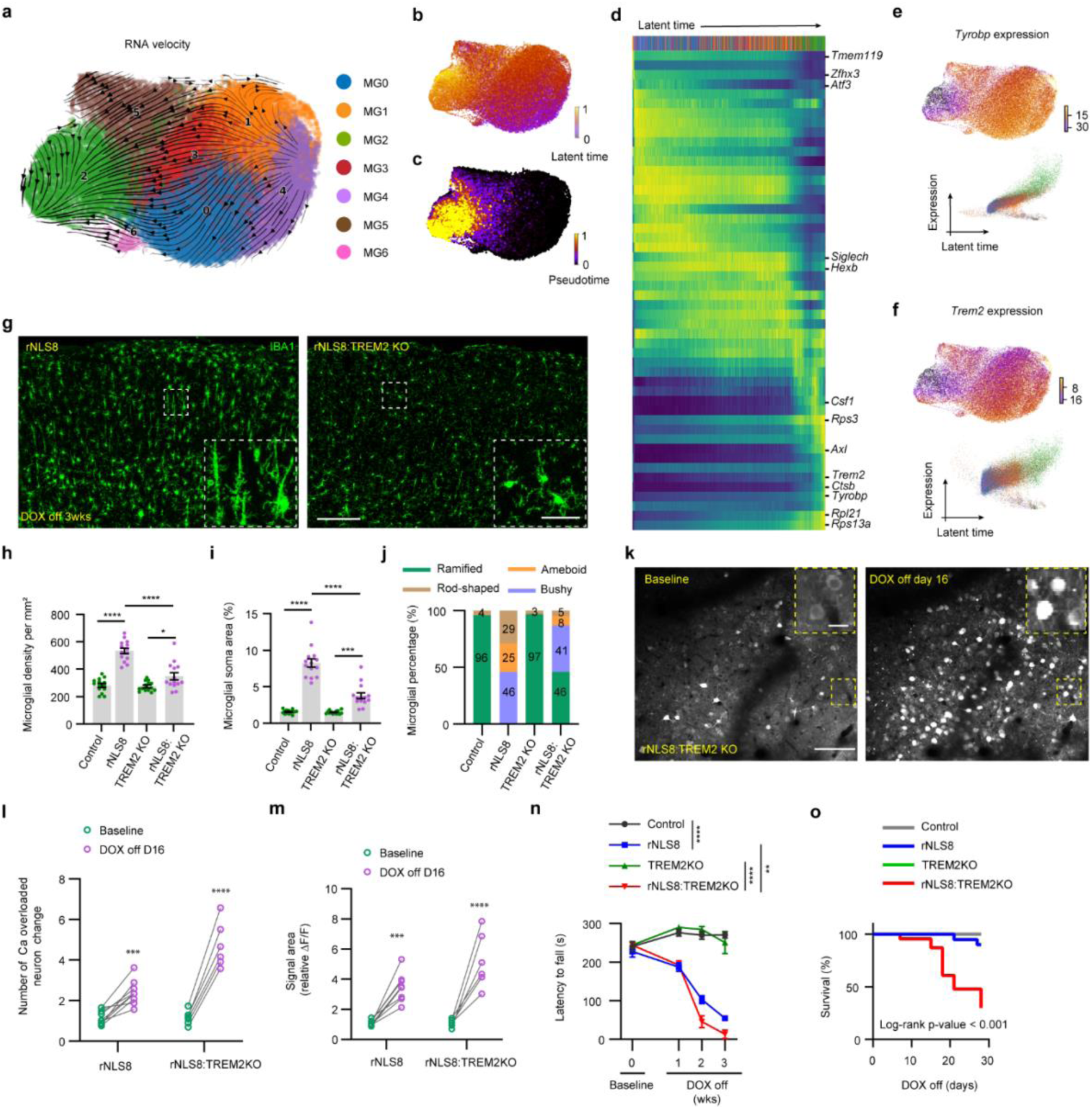
TREM2/DAP12 axis regulates the transition of rod-shaped microglia and neuronal hyperactivity. **a,** RNA velocity derived from the dynamical model for microglia subclusters is visualized as streamlines in a UMAP-based embedding with scVelo. The dynamic model accurately delineates the trajectory of mRNA transcription states among microglia clusters. **b,c,** The scVelo’s latent time (**b**) and pseudo time (**c**) are based on the transcriptional dynamics and visualized in UMAP plots. **d,** Heatmap of the 100 selected highly dynamic genes along latent time shows a clear cascade of transcription. **e-f,** The expression plots (top) and referred expression levels along latent time (bottom) of *Tyrobp* (**e**) and *Trem2* (**f**). **g,** Representative images of microglia (IBA1, green) in M1 of control and rNLS8 mice at 3 weeks post-DOX diet removal. Scale bar, 100 µm. Inset shows microglia at higher magnification as indicated by the area in the dotted white box in the motor cortex. Scale bar, 20 µm. **h, i,** Quantification of microglial density (**h**) and soma size (**i)** in M1 of the indicated groups at 3 weeks post-DOX diet removal (*n* =14 per group). **j**, Percentage of microglia with different morphologies in the indicated groups at 3 weeks post-DOX diet removal. **k**, Representative images of layer 2/3 neuronal calcium activity across experimental phases in rNLS8:TREM2 KO mice during baseline and disease progression. Scale bars, 100 μm. Insets show neuronal soma at higher magnification as indicated by the area in dotted yellow boxes. Scale bar, 10 µm. **l,** Quantification of the number of Ca-overloaded neurons of the indicated groups at baseline and 16 days post-DOX diet removal. **m,** Quantification of ΔF/F signal area (ratio) of neuronal Ca activity in the indicated groups at baseline and 16 days post-DOX diet removal. **n,** Average latency to fall during rotarod tests in the indicated groups during baseline and disease progression (*n* = 25 per group). **o,** Kaplan–Meier survival curves showing the percentage of mice alive at each postnatal day up to 30 days post-DOX diet removal in the indicated groups (*n* = 25 per group). Statistical analysis, one-way ANOVA followed by Tukey’s post hoc test (**h** and **i**), two-tailed paired t-test (**l** and **m**), or two-way ANOVA followed by Sidak’s post hoc test (**n**). Survival curves were analyzed using a log-rank (Mantel–Cox) test (**o**). Error bars, mean ± s.e.m. NS = not significant, \**P* < 0.05; \*\**P* < 0.01; *** *P* < 0.001; **** *P* < 0.0001.

To validate this hypothesis, we crossed rNLS8 mice with TREM2 knockout (KO) mice and evaluated the microglial response during disease progression. Indeed, TREM2 deficiency significantly dampened the microglial response at three weeks post-DOX diet removal, characterized by decreased microglial density and reduced soma size (**Fig. 6g-i**). Morphological analysis of microglia indicated that TREM2 deficiency significantly reduced the number of rod-shaped microglia in rNLS8 mice (**Fig. 6j**).

To evaluate the effect of rod-shaped microglia abolishment on neuronal activity, we conducted longitudinal calcium imaging in rNLS8:TREM2 KO mice. Notably, we observed a significant increase in calcium overload at 16 days post-DOX diet removal in rNLS8:TREM2 KO mice compared with rNLS8 mice, characterized by the nuclear distribution of calcium signals (**Fig. 6k,l and Supplementary Videos 8 and 9**). Compared with ring-shaped neurons, the analysis of the ΔF/F calcium traces indicated that neurons experiencing calcium overload exhibited prolonged duration and elevated amplitude of calcium events (**Extended Data Fig. 10e-g**). Consequently, the area of the calcium signal showed a dramatic increase in rNLS8:TREM2 KO mice (412 ± 19%) compared with those in rNLS8 mice (228 ± 49%) (**Fig. 6m**). In addition, behavioral tests revealed more severe motor deficits and a lower survival rate in the rNLS8:TREM2 KO mice compared to rNLS8 mice, suggesting a neuroprotective effect of the TREM2-mediated microglial response during disease progression (**Fig. 6n,o**). Altogether, these data indicate that the TREM2/DAP12 axis plays a crucial role in mediating the transformation of rod-shaped microglia. The absence of rod-shaped microglia due to TREM2 deficiency aggravates the overall disease outcome, highlighting TREM2-mediated neuroprotective function of rod-shaped microglia.

## Discussion

Microglial activation is one of the hallmarks of ALS pathology ^16,23-26^. However, the exact role of microglia in ALS is not fully understood. Previous studies suggest that microglia exhibit a dual function in ALS, encompassing both neuroprotective and neurotoxic effects ^16,27-29^. This may be attributed to differences in disease stages and models used between studies. Growing evidence indicates that microglia play a crucial role in monitoring and regulating neuronal activity under both normal and disease conditions ^9,10,30^. Yet, the response of microglia to neuronal hyperexcitability in ALS is understudied. Using silicon probes and chronic *in vivo* two-photon calcium imaging, we recapitulated neuronal hyperactivity in the motor cortex of an ALS mouse model in the early stages of disease. Single-cell and spatial RNA sequencing identified microglia, particularly a unique subpopulation of rod-shaped microglia, as the main responders. These rod-shaped microglia have a distinctive transcriptional profile and are pivotal in engaging directly with dendrites and modulating excitatory inputs through the engulfment of excitatory synapses. Rod-shaped microglia formation is heavily reliant on the TREM2/DAP12 axis. TREM2 deficiency led to a marked decrease in rod-shaped microglia, resulting in increased neuronal activity, more severe motor impairments, and lower survival rates in rNLS8 mice. Our findings highlight the novel neuroprotective effects of rod-shaped microglia in the context of TDP-43 neurodegeneration.

Cortical hyperexcitability in the early stages of ALS is believed to be accompanied by progressive and selective degeneration of the motor cortex. The upper motor neurons (the largest pyramidal or Betz cells) primarily found in layer 5, receive excitatory input mainly from pyramidal neurons in the upper layers (mainly layer 2/3), and inhibitory input from local inhibitory neurons.^31^. Our study provides comprehensive characterization of motor cortical hyperexcitability in the rNLS8 mouse model with high spatial and temporal resolution. While previous studies have used electrophysiological and neuroimaging techniques to explore changes in motor neuron excitability in ALS mouse models ^3,32,33,34^, none have conducted longitudinal studies to directly monitor neuronal activity throughout the disease progression with such detailed spatial and temporal resolution. Our findings reveal early hyperactivity in pyramidal neurons across both the upper and lower motor cortical layers. Hyperactivity is most closely related to changes in the firing properties of pyramidal neurons, rather than reductions of inhibitory neuron output at this stage. Following this hyperactive phase, we note a decrease in both the firing rate of pyramidal neurons as well as inhibitory neurons, indicating a disruption of the motor cortical microcircuits. Further research is needed to understand the progression of motor neuron circuit changes in later stages of ALS.

One of the major strengths of our study is the detailed profiling of transcriptional changes associated with neuronal hyperactivity in rNLS8 mice through the integration of spRNA-seq and scRNA-seq. Specifically, our study revealed a unique subpopulation of microglia, rod-shaped microglia, during the short time window of initial disease progression. Rod-shaped microglia were reported in pathological conditions such as neurodegenerative diseases. For example, pathology study from AD patients revealed rod-shaped microglia in close proximity to senile plaques ^35^. They were also observed in the CA1 regions of AD hippocampus, which were systematic overlapped with paired helical filaments (PHF1) positive neurites ^36^. In PD patients, rod-shaped microglia have been observed in the substantia nigra area and were close proximity to degenerating dopaminergic neurons ^37^. In HD patients, rod-shaped microglia were one of the predominant reactive microglia in the cortex. The processes of rod-shaped microglia were aligned along the dendrites and soma of pyramidal neurons ^38^. However, the formation and the function of rod-shaped microglia in neurodegeneration are largely unknown. In addition, rod-shaped microglia have not been reported in the context of ALS/FTD previously. Here we identified this unique type of microglia in the motor cortex of an ALS transgenic mouse model, rNLS8 mice. More interestingly, we found rod-shaped microglia in postmortem tissue samples from ALS patients, suggesting their clinical relevance in ALS pathophysiology. There is no specific marker to distinguish rod-shaped microglia from other microglial subpopulations so far. In this study, we found that 60% of rod-shaped microglia could be identified by Galectin-3 (a member of the lectin family), whereas control microglia exhibited very low expression. This observation is supported by prior findings from ALS patients and mouse models, exhibiting increased levels of Galectin-3 in activated microglia ^39,40^, as well as in plasma and cerebrospinal fluid ^41-44^. The function of microglial Galectine-3 in AD was reported to act as an endogenous ligand for TREM2 and involved in microglial function ^45^. However, limited studies have suggested that Galectine-3 might play a potential role in microglial activation and phagocytosis in ALS ^39,46^. Thus, further research is needed to fully understand whether and how Galectin-3 regulate the formation and function of rod-shaped microglia in the context of ALS.

We were intrigued to observe that the majority of rod-shaped microglia align along neuronal dendrites, particularly the apical dendrites of pyramidal neurons in the rNLS8 mice. This is consistent with previous studies indicating that neuronal processes guide rod-shaped microglia positioning ^47,48^. Gene enrichment analysis further points towards a cell-cell interaction mechanism. Thus, we propose that rod-shaped microglia are able to sense and regulate neuronal function via direct interaction with neuronal dendrites in the cortex of rNLS8 mice. Growing evidence suggests that microglia engage in neuronal circuits in healthy and neurological disorders ^11,49-51^. Established mechanisms through which microglia sense neuronal activity include ATP/purinergic signaling ^11,49-51^, microglia Ca^2+^ signaling ^52^, fractalkine signaling ^53^, neuromodulators ^54,55^, complement signals ^56^, and others. Considering the neuronal hyperactivity is associated with rod-shaped microglia formation in rNLS8 mice, it will be important to identify the signaling pathways that enable rod-shaped microglia to sense neuronal hyperactivity. In addition, the causal relationship between rod-shaped microglia and neuronal hyperactivity needs to be further addressed.

Microglia-mediated synaptic pruning, a well-documented phenomenon across various neurodegenerative disease models, involves mechanisms such as completement ^57^, CX3CR1 ^58^, TREM2 ^59,60^, and others. In the context of ALS, it has been reported that C9orf72-depleted microglia exhibit enhanced cortical synaptic pruning ^61^. Here we found that rod-shaped microglia have a high capacity for phagocytosing excitatory synaptic materials, suggesting their potential role in regulating neuronal activity in rNLS8 mouse model. Our receptor-ligand–based cellular communication analysis uncovered several putative candidate molecules that may facilitate their interaction. For example, cell adhesion and phagocytosis related ligand-receptor pairs. Future research is necessary to validate whether these receptor-ligand pairs are involved in the interaction between rod-shaped microglia and neuronal dendrite, and its potential function in synapse elimination and neurodegeneration.

We found that rod-shaped microglia peaked at 3 weeks after disease progression in rNLS8 mice. In addition, they exhibited a specific distribution across cortical layer 2 to 4. Thus, rod-shaped microglia represent a transitional state in spatiotemporal manner. The key question is how homeostatic microglia transformed into rod-shaped microglia. To this end, we discovered that the TREM2/DAP12 axis plays a crucial role in the emergence of rod-shaped microglia, as determined by trajectory analysis. In the central nervous system (CNS), TREM2 is exclusively expressed by microglia and is involved in various functions such as proliferation, activation, and phagocytosis ^29,62,63^. The role of TREM2 in modulating microglial status in AD is well-documented, facilitating the transition to an activated state and the induction of lipid metabolism and phagocytic pathways ^18^. Our previous studies have found that TREM2 interacts with TDP-43 and mediates microglial phagocytosis of TDP-43 that is important for disease recovery^16^. Here we observed that TREM2 deficiency significantly reduced the formation of rod-shaped microglia while increased the neuronal activity. In addition, TREM2 deficiency worsened motor deficits and lower survival rates, suggesting potential neuroprotective function of rod-shaped microglia at early phase of disease progression. Known TREM2 ligands, such as phosphatidylserine (PS), TDP-43, and damaged lipids ^16,63,64^, may be produced or released by hyperactive neurons and serve as cues for TREM2. Future research should aim to ascertain how TREM2 detects signals from hyperactive neurons, thereby regulating the transformation of rod-shaped microglia.

In sum, our study reported the neuronal hyperactivity and associated spatiotemporal formation of rod-shaped microglia in rNLS8 mice. These findings suggest that rod-shaped microglia may exert neuroprotective function through elimination of the excitatory synaptic inputs, highlighting the therapeutic potential of microglia in early ALS treatment.

## Methods

### Mice

The rNLS8 mouse line was crossed by the NEFH-tTA line 8 (JAX, No. 025397) and tetO-hTDP-43-ΔNLS line 4 (JAX, No. 014650). Mice hemizygous for both NEFH-tTA and tetO-hTDP-43-ΔNLS were used as diseased mice. Mice hemizygous for only NEFH-tTA or tetO-hTDP-43-ΔNLS were used as control mice. Thy1-YFP-H (JAX, No. 003782) and *CX_3_CR-1^GFP^* knock-in/knock-out (JAX: 005582) mice were purchased from Jackson Labs. The TREM2 knockout (KO) mouse strain was generously provided by Dr. Marco Colonna at the Washington University School of Medicine, St. Louis. Thy1-YFP-H and TREM2 KO mice were crossed with rNLS8 mice. rNLS8 mice were maintained on Dox chow (200 mg/kg, Bio-Serv #3888) to prevent hTDP-43ΔNLS expression. Experimental mice were switched to Dox-free chow to induce hTDP-43ΔNLS expression. Mouse lines were housed with littermates with free access to food and water on a 12-hour light/dark cycle, and the room temperature is 20-22°C with 55% humidity. All animal procedures, including husbandry, were performed under the guidelines set forth by Mayo Clinic Institutional Animal Care and Use Committee. Both sexes were used in the experiments.

### Human samples

Post-mortem brain samples were dissected from the frozen brains of 31 ALS cases (age 61.32 ± 9.70 years, mean ± s.d.) and 25 controls (age 59.09 ± 12.73 years) from the Brain Bank for Neurodegenerative Disorders at Mayo Clinic Jacksonville. Brainstem-type Lewy body disease (BLBD) cases with minimal p62 immunoreactivity were identified as controls for motor cortex pathology. The study was approved by Mayo Biospecimen Committee. For clinic pathological studies, cases were included only if they had good quality medical documentation and there was diagnostic concurrence of at least minimum of two neurologists.

### *In vivo* multichannel silicon probes recording

#### Silicon probe implantation surgery

All recordings were performed using A1x32-Poly2-3mm-50 s-177-CM32 silicon probes (177 μm^2^ site surface area, 2-column honeycomb site geometry with 50 μm center-to-center span, 15 μm shank thickness) with a CM32 connector (NeuroNexus Technologies). Probe implantation surgery is performed based on the protocol provided by the NeuroNexus website. Specifically, animals were placed on a stereotaxic frame and under isoflurane anesthesia (4% induction, 1.5–2.5% maintenance). A circular craniotomy (1 mm diameter) was drilled to accommodate the probe shanks in the mouse primary motor cortex at the following stereotactic coordinates: AP, - 1.4 mm from bregma; ML, 1.5 mm; DV, - 0.8 mm from the brain surface. Additionally, three small holes (1 mm diameter) were then drilled for stainless steel screws: two over both sensory cortices and one over the contralateral site of the implant, which were later used as a ground reference. Screws (4 mm long, 0.86 mm diameter) were carefully advanced into the drilled holes. After removal of the dura in the craniotomy, probes were lowered into the surface of the cortex and inserted using an automatic or manual micromanipulator to the desired depth (0.8mm) at a rate of 1 mm/min. The ground and reference wires were wrapped tightly around the bone screws before the probe and wires were secured using dental cement and a secure head cap was created. Mice were kept warm and accelerated recovery from anesthesia, then single-housed to protect the electrode.

#### Data collection and analysis

Ten days following the procedure, mice were acclimated to the recording environment by placing them in a recording box with their electrodes connected to the recording adaptor for one hour daily. This habituation period lasted approximately 3 days, during which the mice were allowed to move freely within the recording box. During the recording sessions, mice were given 30 minutes to acclimate to the environment with their electrodes connected to the adapter before the recording commenced. Recordings were carried out weekly for a duration of 30 minutes each, covering both the baseline and disease progression periods over four weeks. The head stage was connected to a Zeus recording system (Zeus, Bio-Signal Technologies, McKinney, TX, USA) to capture peak potentials, using a sampling frequency of 3 kHz. Spike signals were band-pass filtered online at 300-7,000 Hz. Spike waveforms and timestamps were saved in Plexon data (*.plx) files. Spikes were sorted using the valley seek method with Offline Sorter software (Plexon), and any indistinct waveforms were manually removed in a three-dimensional feature space. The sorted units were analyzed with Neuro Explorer 5.0 to generate graphs and charts detailing the firing rates and timestamps of the neurons, which were then exported for further analysis in Microsoft Excel.

Data analysis was performed as previously described ^65-67^. Units were classified as either excitatory or inhibitory neurons based on their spike width (trough-to-peak duration): units with a spike width greater than 0.4 ms were classified as excitatory neurons, while those with a spike width less than 0.4 ms were classified as inhibitory neurons. Additionally, the mean interspike interval (ISI) was used to further distinguish between excitatory and inhibitory neurons.

Inhibitory neurons exhibited a shorter ISI compared to excitatory neurons, with putative interneurons having a mean ISI of 2 ms or less. The firing frequency of neuronal activity was analyzed and normalized to the baseline.

#### *Ex vivo* electrophysiological recordings

Brain slices containing the motor cortex were prepared for electrophysiological recordings as previously described ^68^. Mice aged P70 were deeply anesthetized using isoflurane. Following decapitation, the brain was quickly removed and submerged in an ice-cold slice solution, which was oxygenated with 95% O_2_ and 5% CO_2_. This solution contained the following components (in mM): 2.41 KCl, 1.22 NaH_2_PO_4_·2H_2_O, 25 NaHCO_3_, 0.4 ascorbic acid, 2 sodium pyruvate, 0.5 CaCl2·2H2O, 3.49 MgCl2·6H2O, and 240 sucrose; it was adjusted to a pH of 7.4 ± 0.5 using HCl. Coronal slices of 300 μm thickness were sectioned in the cold slice solution using a Leica 2000 vibratome. These slices were then allowed to recover in an oxygenated slice solution at 34°C for 13 minutes, followed by maintenance in an incubation chamber filled with oxygenated artificial cerebrospinal fluid (ACSF) at room temperature for 30 minutes before the recordings. The ACSF composition was as follows (in mM): 117 NaCl, 3.6 KCl, 1.2 NaH_2_PO_4_·2H_2_O, 25 NaHCO_3_, 0.4 ascorbic acid, 2 sodium pyruvate, 2.5 CaCl_2_·2H_2_O, 1.2 MgCl_2_·6H_2_O, and 11 glucose, adjusted to a pH of 7.4 ± 0.5 with HCl. All chemicals were sourced from Sigma.

Whole-cell recordings were performed at room temperature in the oxygenated ACSF solution. The brain slices were observed under a fixed upright microscope (Scientifica) equipped with a water immersion lens (Olympus, 40x/0.8 W). Patch pipettes were fashioned from borosilicate glass capillary tubes using a horizontal pipette puller (P-97, Sutter Instruments). These pipettes were filled with a potassium-based internal solution for recording active potentials in current-clamp modes, with the neuronal membrane potential held at -70 mV. The composition of the internal solution was (in mM): 135 potassium gluconate, 0.5 CaCl_2_·2H_2_O, 2 MgCl_2_·6H_2_O, 5 KCl, 5 EGTA, and 5 HEPES; the pH was adjusted to 7.3 ± 0.5 using KOH, achieving an osmolarity of 290-300 mOsm. The resistance of the patch pipettes was between 5-7 MΩ.

Data collection was facilitated by a Multiclamp 700B amplifier (Axon Instruments), with the signal filtered at 2 kHz and digitized at 4 kHz using a Digidata 1440A converter (Axon Instruments). Analysis of the recorded data was conducted using Clampfit 10.3 (Axon Instruments).

### Chronic *in vivo* Ca imaging

#### Stereotaxic delivery of AAV

Adult rNLS8 or control mice at the age of 2 months were anesthetized by isoflurane (4% for induction and 2.5% for maintenance). Subsequently, 250 nL of pENN.AAV9.CaMKII.GCaMP6s.WPRE.SV40 (concentration of 1.0E+12 viral genomes/mL, Addgene, 107790) was injected into the right primary motor cortex to specifically target layer 2&3 neurons. The coordinates for the injections were as follows: 1.4 mm anterior to bregma, 1.5 mm lateral to the midline, 0.3 mm ventral to the dura, with bregma set at zero. These microinjections were administered at a rate of 40 nL/min, utilizing a glass pipette and an automated stereotaxic injection apparatus (World Precision Instruments Inc). To ensure optimal diffusion of the viral vector, the microsyringe was maintained in position for an additional 10 minutes following each injection.

#### Cranial window surgery

Under isoflurane anesthesia (4% for induction and 2.5% for maintenance), a circular craniotomy of 4 mm in diameter was created over the motor cortex using a high-speed dental drill. A circular glass coverslip, also 4 mm in diameter (Warner), was then securely placed over the craniotomy site. Additionally, a four-point headbar (NeuroTar) was affixed atop the window using dental cement. To manage pain, a mild NSAID solution (0.2 mg/mL Ibuprofen) was administered in the mice home cage for 72 hours both before and after the surgery. For mice undergoing AAV injections, the cranial windows were implanted concurrently with the injection process.

#### Data collection

Ca imaging in awake animals was conducted using a multiphoton microscope, equipped with galvanometer scanning mirrors (Scientifica). To visualize GCaMP6s fluorescence, we employed a Mai-Tai DeepSee laser (Spectra Physics), tuning it to 920 nm and maintaining the output below 55 mW for layer 2/3 imaging. The GCaMP6s signals were then filtered through a 520/15 filter (Chroma). Imaging was performed at a frame rate of 1 Hz, with a resolution of 512 × 512 pixels and a zoom factor of 450 × 450 µm, using a 16x water-immersion objective (Nikon, 16x; NA: 0.8). Before data collection, mice were trained to move on an air-lifted platform (NeuroTar) while being head-fixed under a two-photon (2P) objective. This training was conducted for 30 minutes per day over a period of 7 days prior to the experiment. Chronic imaging sessions were scheduled 3-4 weeks post-surgery in mice that exhibited a clear window. In all studies, we allowed the mice a 10-minute acclimation period in the head restraint before beginning imaging. For each time point during the imaging sessions, a 15-minute video was recorded. Mouse locomotion was recorded during *in vivo* calcium imaging using the mobile home cage magnetic tracking system (Neurotar).

#### Calcium image processing and analysis

Time series data of neuronal calcium activity were subjected to motion correction using the TurboReg plugin in ImageJ. Subsequently, an average intensity image was generated for the selection of Regions of Interest (ROIs). For identifying neuronal somata, visually discernible cell bodies were manually selected as ROIs using ImageJ. Once the ROIs were established, the multi-measure tool was employed to acquire mean intensity values (F) for each ROI. The relative percentage change in fluorescence of the calcium signals was calculated using the formula ΔF/F = (F − F0)/F0. Here, the baseline F0 was defined as the lower 25th percentile value of the initial 200-ms period of the recording. In instances of heightened activity, the baseline was alternatively determined by identifying the lower 25th percentile value in a user-selected, 200-frame period exhibiting minimal wave activity. A calcium transient was considered to have occurred when the ΔF/F ratio exceeded a threshold threefold greater than the standard deviation of the baseline. The signal area was quantified as the sum of all ΔF/F values surpassing this threshold over the entire duration of the recording. The maximum amplitude was calculated as the highest peak value among all the transients.

### Spatial RNA sequencing (spRNA-seq)

#### 10X Visium preparation and sequencing

Mice were anesthetized with isoflurane (5% in O_2_) and then intracardially perfused with 40 ml cold PBS, followed by 40 ml cold 4% paraformaldehyde (PFA). Brains were removed and post-fixed in 4% PFA for an additional 6-8 hours or overnight in RT. Samples were then raised by 1X PBS and then processed by the following steps using the automatic tissue processing machine: 70% Alc (1 hr), 80% Alc (1 hr), 95% Alc (30 min), 95% Alc (30 min), 95% Alc (30 min), 100% Alc (30 min), 100% Alc (30 min), 100% Alc (45 min), Xylene (60 min), Xylene (60 min), Paraffin (45 min 60° C), Paraffin (45 min 60° C), Paraffin (60 min 60° C), Paraffin (60 min 60° C). The tissues were then embedded in paraffin to create a paraffin block. The paraffin blocks were then cut using a microtome to generate thin sections of tissue for RNA quality assessment and 10X Visium preparation. After RNA quality assessment (DV200 > 50%), the FFPE tissue block was then sectioned by a microtome at 5µm to generate appropriately sized sections for Visium slides (app. Bregma 1.54 mm; Allen brain reference atlas coronal section 18). The slides were then placed in a slide drying rack and incubate for 3 h in an oven at 42°C (10X Genomics, CG000408). After overnight drying at room temperature, the slices were proceeded to deparaffinization, decrosslinking and immunofluorescence staining protocols according to the manufacturer’s protocol with recommended reagents (10X Genomics, 1000339 and 1000251; CG000410). After tissue imaging, the slides were proceeded immediately to Visium Spatial Gene Expression based on User Guide (10X Genomics, CG000407), including probe hybridization, probe ligation, probe release & extension. Library generation commenced with the amplification of eluted probes using real-time qPCR to ascertain the optimal number of amplification cycles. The qPCR protocol commenced with an initial denaturation step lasting 3 minutes at 98°C, succeeded by 25 cycles, each comprising a 5-second duration at 98°C and a 30-second duration at 63°C. The requisite number of PCR cycles for library amplification was established based on the qPCR outcomes, employing 16-19 cycles for the amplification of the entire set of eluted probes, utilizing a 10X dual index kit. Subsequently, the amplified libraries were purified with SPRI select beads (Beckman Coulter). Quantification of the final library was conducted using a TapeStation 4200 D1000 Screen Tape (Agilent) and Qubit (Invitrogen). Sequencing was performed employing paired-end reads of 101 bp, utilizing either the NextSeq2000 P2 or NovaSeq6000 SP.

### spRNA-seq quality control, integration, clustering, and differential expression analysis

Quality control was performed on each sample to remove low-quality spots and lowly expressed genes. Spots with low total UMI counts, low numbers of expressed genes, high mitochondria concentration, or high hemoglobin concentration were removed. Genes detected in fewer than 10 spots were also removed. The data were then normalized using SCTransform v2 and integrated using the Harmony algorithm ^69^. Gene-expression-based clusters were generated for each sample, with dimension reduction and clustering implemented using the Seurat R package ^70^. These clusters were then annotated based on pathology review of the corresponding slides within Loupe Browser. Spots that passed the initial quality control but were found to be outside the tissue area during pathology review were removed. Differential gene expression analysis was performed to identify differentially expressed genes between each cluster and to identify differences in the disease phenotype within each cluster. The FindMarkers function from the Seurat R package was used to perform the DESeq2 test. Genes with an absolute value of log2(fold change) greater than 0.5 and a Bonferroni-adjusted p-value less than 0.05 were considered significant. Gene Ontology (GO) analysis was performed with the clusterProfiler R package ^71^. Upregulated pathways were determined by analyzing sets of genes significantly upregulated in the disease samples, and additional pathways were determined by analyzing the full set of significantly differentially expressed genes. The Benjamini-Hochberg method was used to control the false discovery rate.

### Single cell RNA sequencing

#### Single-cell preparation and sequencing

Single cell preparation was performed as previously described ^72^. Mice were transcardially perfused with cold PBS and cortical regions were quickly dissected and enzymatically digested using the Neural Tissue Dissociation Kit P (Miltenyi Biotec, 130-092-628) in the gentleMACS Octo Dissociator with heaters (Miltenyi Biotec) using program 37C_NTDK_1. The myelin was then removed by magnetic bead separation using Myelin Removal Beads II (Miltenyi Biotec, 130-096-733) followed by the red blood cell lysis (Miltenyi Biotec, 130-094-183). Cell suspension was then followed by dead cell removal (Miltenyi Biotec 130-090-101;). The final cell pellet was suspended in Ca/Mg-free PBS with 0.5% BSA, and immediately submitted to the Genome Analysis Core for Single Cell partitioning. The cells were counted and measured for viability using the Vi-Cell XR Cell Viability Analyzer (Beckman-Coulter). The barcoded Gel Beads were thawed from -80 °C and the cDNA master mix was prepared according to the manufacture’s instruction for Chromium Next GEM Single Cell 3’ Kit v3.1 (10x Genomics). Based on the desired number of cells to be captured for each sample, a volume of live cells was mixed with the cDNA master mix. A per sample concentration of 500,000 cells per milliliter or better is required for the standard targeted cell recovery of up to 10,000 cells. The stock concentration requirements would not change for higher cell recovery numbers. The cell suspension and master mix, thawed Gel Beads and partitioning oil were added to a Chromium Next GEM G chip. The filled chip was loaded into the Chromium Controller, where each sample was processed and the individual cells within the sample were partitioned into uniquely labeled GEMs (Gel Beads-In-Emulsion). The GEMs were collected from the chip and taken to the bench for reverse transcription, GEM dissolution, and cDNA clean-up. The resulting cDNA contains a pool of uniquely barcoded molecules. A portion of the cleaned and measured pooled cDNA continues to library construction, where standard Illumina sequencing primers and a unique sample index (Dual Index Kit TT-Set A; 10x Genomics) were added to each cDNA pool, creating gene expression libraries. All cDNA pools and resulting libraries are measured using Qubit High Sensitivity assays (Thermo Fisher Scientific) and Agilent Bioanalyzer High Sensitivity chips (Agilent). Libraries are sequenced at 50,000 fragment reads per cell following Illumina’s standard protocol using the Illumina NovaSeq™ 6000 S4 flow cell. S4 flow cells are be sequenced as 100 × 2 paired end reads using NovaSeq S4 sequencing kit and NovaSeq Control Software v1.8.0. Base-calling is performed using Illumina’s RTA version 3.4.4.

### scRNA-seq quality control, integration, clustering, ligand-receptor interaction and differential expression analysis

Single-cell sequence preprocessing was performed using the standard 10x Genomics Cell Ranger Single Cell Software Suite. Raw reads were aligned to the hg38 reference genome, and UMI (unique molecular identifier) counting was performed using Cell Ranger v7.1.0 with default parameters. Seurat v5 was used for all subsequent analyses ^73^. Genes not detected in at least three single cells were excluded. Cells with fewer than 500 unique genes detected were also excluded to remove potential low-quality cells. Mitochondrial quality control metrics were calculated using the ‘PercentageFeatureSet’ function to filter out cells with >20% mitochondrial counts, thereby avoiding low-quality and dying cells. Normalization was performed with the ‘LogNormalize’ function in Seurat, followed by log transformation for downstream analysis. The ‘FindVariableFeatures’ function was then used to identify a subset of highly variable features for each sample, highlighting biological signals for future analyses. For integration analysis and batch effect removal, the Harmony R package (harmony_1.1.0) was utilized ^69^. Principal component analysis (PCA), a dimensionality reduction technique, was conducted to determine the dataset’s dimensionality, and UMAP was chosen to visualize the combined dataset. The ‘FindNeighbors’ and ‘FindClusters’ functions were used to cluster cells. Cluster-specific markers conserved across conditions were identified using the ‘FindConservedMarkers’ function, and clusters were assigned to known cell types based on these markers.

After manual annotation of all cell clusters as specific cell types, comparative analysis was performed using the MAST ^74^ model to identify differentially expressed genes (DEGs) induced by different conditions. GO analysis was performed using the clusterProfiler package ^71^. DEGs identified from the comparative analysis were used as input for the GO enrichment analysis. The ‘enrichGO’ function was applied to categorize DEGs into biological processes, molecular functions, and cellular components. The results were visualized using the ‘dotplot’ function to highlight significant GO terms.

The ligand-receptor interaction analysis was performed using NicheNet v2 ^20^. We utilized the Seurat v5 object provided by the Department of Quantitative Health Sciences at Mayo Clinic. Our methodology closely followed the vignettes available on GitHub from saeyslab/nichenetr, specifically “Perform NicheNet analysis starting from a Seurat object” and “Seurat Wrapper + Circos visualization.” First, we loaded the ligand-receptor network, ligand-target matrix, and weighted networks for mice from https://zenodo.org/records/7074291. From our original Seurat object, we created a subset Seurat object that included only cells classified as “Microglia” and “Neuron,” with “Microglia” also encompassing the subset labeled as “Mitotic microglia.” After re-normalizing the raw counts of the subsetted object, we ran ‘nichenet_seuratobj_aggregatè with the sender population set to “Neuron” and the receiver population set to “Microglia.” We configured this function to return the top 30 ligands and top 200 gene targets. Finally, we used the output from this function to create a Circos plot, which facilitated better visualization of the identified interactions.

For RNA Velocity analysis, the Velocyto pipeline was used to quantify spliced and unspliced RNA counts from the single-cell RNA sequencing data ^21^ and generate loom files. Gene-specific velocities were then computed using the scVelo package ^22^. The ‘scvelo.pp.filter_and_normalize’ function was used for pre-processing, including filtering and normalization of the data. The function ‘scvelo.pp.moments’ is applied to capture transcriptional dynamics and RNA velocity vectors were estimated using the ‘scvelo.tl.velocity’ function. Finally, the velocities were projected onto the existing UMAP embedding using the ‘scvelo.tl.velocity_graph’ and ‘scvelo.pl.velocity_embedding’ functions to visualize the dynamic changes in gene expression over time. The latent time was calculated using the ‘scv.tl.latent_time’ function, which helps in identifying the progression of cells through different states. The results were visualized with ‘scv.pl.heatmap’, which displays the expression dynamics of top genes over the inferred latent time, and ‘scv.pl.scatter’, which plots the latent time on the UMAP embedding, allowing for visualization of temporal progression across the cell population.

#### Immunofluorescence staining

Mice were anesthetized with isoflurane (5% in O_2_) and then intracardially perfused with 40 ml cold PBS, followed by 40 ml cold 4% paraformaldehyde (PFA). Brains were removed and post-fixed in 4% PFA for an additional 6 hours, then transferred to 30% sucrose in PBS for three days. Tissue was then embedded in OCT and frozen before being cryosectioned using a Leica Cryostat at 20 μm. For immunofluorescence staining, the sections were blocked for 60 mins with 10% goat or donkey serum in TBS buffer containing 0.4% Triton X-100 (Sigma), and then incubated overnight at 4 °C with a primary IgG antibody (**Supplementary file 8**). After three washes with TBST, sections were exposed to appropriate secondary antibody (**Supplementary file 8**) for 60 mins at room temperature, washed and mounted and coverslipped with Fluoromount-G (SouthernBiotech). Fluorescent images were captured with a confocal microscope (LSM980, Zen software, Zeiss) in the primary motor cortex. Images were taken from a single Z-plane (1024 x 1024 Pixels). Cells counts, fluorescence signal intensity and area of each cell were quantified using the Analyze Particles function in ImageJ (National Institutes of Health, Bethesda, MD). Volume rendering and visualization were performed using Imaris v.9.2 (Oxford Instruments).

#### IMARIS Rendering

Three-dimensional renderings of rod-shaped microglial interaction with apical dendrites (IBA1 staining in rNLS8:Thy1-YFP-H mouse brain slices) and microglial engulfment of synaptic markers (costaining of IBA1/ VGLUT1/PSD95) were performed in Imaris using representative Z stack images. Z-stack images of full microglia were collected by a 63x objective (oil, NA:1.4), with 2048x2048 pixel rendering and a 0.3 µm step size (LSM980, Zen software, Zeiss). Channels were respectively converted into surfaces using Imaris to preserve structural and spatial integrity. Individual PSD95 and VGLUT1 puncta that were not interacting or engulfed by the IBA1^+^ surface was manually removed. Representative images were taken with the Snapshot function and videos taken with the Animation function.

#### Immunohistochemistry of patient’s tissue

Formalin-fixed brains underwent systematic and standardized sampling with neuropathologic examination by a single board-certified neuropathologist (D.W.D.). Specific brain regions were dissected from the fixed hemibrain and studied for gross and microscopic pathology using the Dickson sampling scheme ^75^. 5 μm thick paraffin-embedded sections were cut. Immunohistochemistry for microglia (IBA-1, rabbit polyclonal, 1:3000, Wako Chemicals, 019-19741) was performed on the motor cortex of 25 ALS and 25 BLBD cases.

#### Nissl staining

Tissue sections were first incubated in 100% ethanol for 6 minutes, followed by a defatting process in xylene for 15 minutes, and then placed back into 100% ethanol for another 10 minutes. After being rinsed with distilled water, the slides were stained with a 0.5% solution of cresyl violet acetate for 15 minutes, followed by another rinse in distilled water. Subsequently, the sections were treated in a differentiation buffer (comprising 0.2% acetic acid in 95% ethanol) for 2 minutes, dehydrated with ethanol and Xylene, and finally mounted with Depex medium. Pyramidal neurons were distinguished by their characteristic triangular shape and the presence of a single, large apical dendrite extending vertically toward the pial surface.

#### Hindlimb clasping

To evaluate the hindlimb clasping behavior, each mouse was carefully lifted from its home cage by suspending it by the tail for a duration of 10 seconds. During this time, the mouse’s hindlimb clasping response was observed and rated on a scale from 0 to 4. A score of 4 was given if the mouse displayed a normal escape response, extending its limbs without any clasping. A score of 3 was assigned if one hindlimb showed partial extension (incomplete splay) along with reduced mobility, yet the toes remained well-spread (normal splay). A score of 2 was given when both hindlimbs showed partial extension and reduced mobility but the toes were spread normally. A score of 1 indicated that both hindlimbs were clasped with curled toes and the mouse exhibited immobility. Finally, a score of 0 was recorded if both forelimbs and hindlimbs were clasped, with crossed, curled toes and the mouse was immobile.

#### Wire hang test

This test measures the force and endurance a mouse exhibits to remain suspended on a wire, countering gravity for as long as possible. The duration of each mouse’s grip is recorded across three attempts, without a set time limit for hanging. The methodology follows previously established protocols (referenced as DMD_M.2.1.004). The initial position of the mice can vary, involving either two or four limbs. The procedure involves the mouse gripping the wire’s midpoint with its front limbs, followed by a gentle adjustment to allow its back paws to grasp the wire, spaced a few centimeters from the front paws. The mouse is then assisted in turning upside-down along the wire’s axis, releasing its tail while it continues to grip with all four paws. A timer is initiated upon tail release, and the period until the mouse releases its grip and falls is noted. Each mouse undergoes three trials in a session, with a 30-second rest between each trial. The average duration from the three trials serves as the primary outcome metric. It’s essential to record the mouse’s body weight either before or after the test, as weight measurements can be essential for longitudinal studies. For tests allowing unlimited hanging time, the influence of body weight on performance can be minimized by calculating the Holding Impulse (s*g) = Body mass (grams) x Hang Time (seconds), offering an adjusted outcome measure.

#### Open field testing

The locomotor activity of the mice was measured in sound-proof, rectangular chambers equipped with fans, infrared lasers, and sensors (Med Associates, St. Albans, VT, USA). Before the tests, the mice were allowed to acclimate to the testing environment for 1 hour. Following this, they were placed in the Open Field chamber for a 10-minute recording period. The primary measure of locomotor function was the total distance traveled by the mice within the chamber, which was recorded and analyzed using the Med-PC software, Version 4.0.

#### Rotarod

The Rotarod performance test was performed to assess the balance and motor coordination of mice. Before the tests, the mice were allowed to acclimate to the testing environment for 1 hour. Following this, the test was conducted on a five-lane Rotarod apparatus (Med Associates Inc.), which began at a speed of 4 revolutions per minute (rpm) and gradually increased to 40 rpm over a span of 5 minutes. Each mouse underwent the test three times, with a rest period of 10 minutes between each trial.

#### Statistics

Statistical details of the experiments, including sample sizes and statistical tests are described in figure legends. Mean values of multiple groups were compared using a one-way ANOVA, followed by a Fisher’s post-hoc test (comparison to baseline), or Tukey’s post-hoc test (comparisons between all groups). Mean values of two groups across time were compared using a two-way ANOVA, followed by either a Sidak’s post-hoc test (comparison between genotype/group at each time point), or a Dunnett’s post-hoc test (comparison of a time point against the baseline). All post-hoc testing accounted for multiple comparisons. Comparison of two groups at a single timepoint was performed using Student’s t-test. Survival curves were analyzed using a log-rank (Mantel–Cox) test. P values < 0.05 were considered as statistically significant. All statistical analysis was performed with GraphPad Prism software (version 10). No statistical methods were used to pre-determine sample sizes, but our samples sizes are comparable to similar studies ^55,76^. Data distribution was assumed to be normal, but this was not formally tested. Mice were grouped according to genotype before they were randomly assigned to the experimental groups. The investigators were blinded to group allocation during data collection and analysis. Animals or samples were excluded from analysis only in the instance of technical failure. No data were excluded for other reasons.

## Data availability

Single cell RNA sequencing and spatial RNA sequencing data have been uploaded to the Sequence Read Archive (SRA) and are publicly available. Other information that supports the findings of this study is available from the corresponding author upon request.

## Acknowledgments

The authors thank Dr. Marco Colonna (Washington University) for providing TREM2 KO mice; Dr. Vanda A. Lennon (Mayo Clinic Rochester) for thoughtful discussion and manuscript editing; Mayo Clinic Rochester Genome Analysis Core for assistance with library preparation, and for single cell RNA sequencing and spatial RNA sequencing. The current study is supported by NIH grants RF1AG082314 and R35NS132326 to L.J.W, and U19AG069701 to D.W.D, and L.J.W.

## Author contributions

M.X. and L.-J.W. developed the research concept, designed the experiments, and wrote the manuscript; M.X. performed most of the experiments and data analysis; A.S.M. performed immunostaining and quantification, and behavioral analysis. A.U. provided knowledge support for two-photon experiments. Y. L. performed electrophysiological experiment and provided knowledge support for silicon probe recording. N.W. provided knowledge support for single cell preparation. M.X., S.Z., N.K.N., Z.C.F., and P.N.P. performed bioinformatics analysis. N.B.G., B.E.O. and D.W.D. provided immunohistochemical staining data from patient tissues. S.Z., A.S.M, and F.Q. performed immunostaining and analysis. J.Z. provided knowledge support for 10X Visium. A.N. provided knowledge support for the study. M.X. and A.S.M. contributed to animal maintenance, genotyping, and tissue collection.

## Competing interests

The authors declare no competing interests.

**Correspondence and requests for materials** should be addressed to Long-Jun Wu

---

**Extended Data Fig 1.**
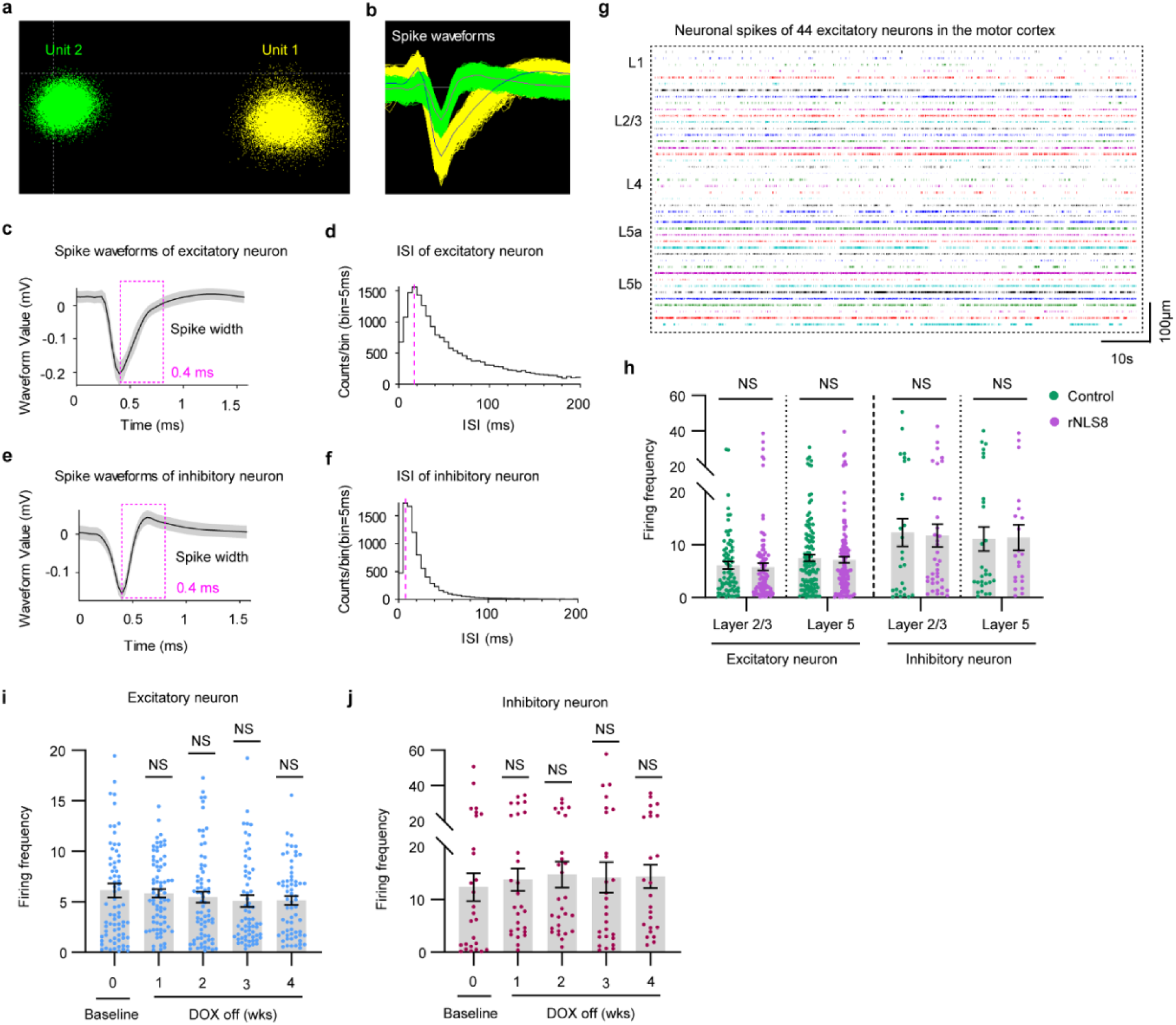
Basic characterization of the silicon probe recording, related to Figure 1. **a,b,** Representative image of two units (**a**, representing two neurons) and their waveform (**b**) recorded by one channel from the silicon probe. **c,d,** Representative images of spike waveforms (**c**) and interspike intervals (ISI) (**d**) of excitatory neurons (pyramidal neurons). **e,f,** Representative images of spike waveforms (**e**) and interspike intervals (ISI) (**f**) of inhibitory neurons (interneurons). **g,** Representative spike rates of isolated pyramidal neurons from all layers of primary motor cortex (M1) during baseline. Scale bars, 100μm (vertical), 10s (horizontal). **h,** Firing frequency of excitatory and inhibitory neurons in layer 2/3 and layer 5 of M1 in control (*n* = 6) and rNLS8 mice (*n* = 7) during baseline. **i,j**, Firing frequency of excitatory (**i**) and inhibitory (**j**) neurons in layer 2/3 of M1 in control mice during baseline and DOX off period (*n* = 6 mice). Statistical analysis, two-tailed unpaired Student’s t-test (**h**), or one-way ANOVA followed by Dunnett’s post hoc comparison to baseline (**i,j**). Error bars, mean ± s.e.m. NS = not significant, \**P* < 0.05; \*\**P* < 0.01; \*\*\**P* < 0.001; \*\*\*\**P* < 0.0001.

**Extended Data Fig 2.**
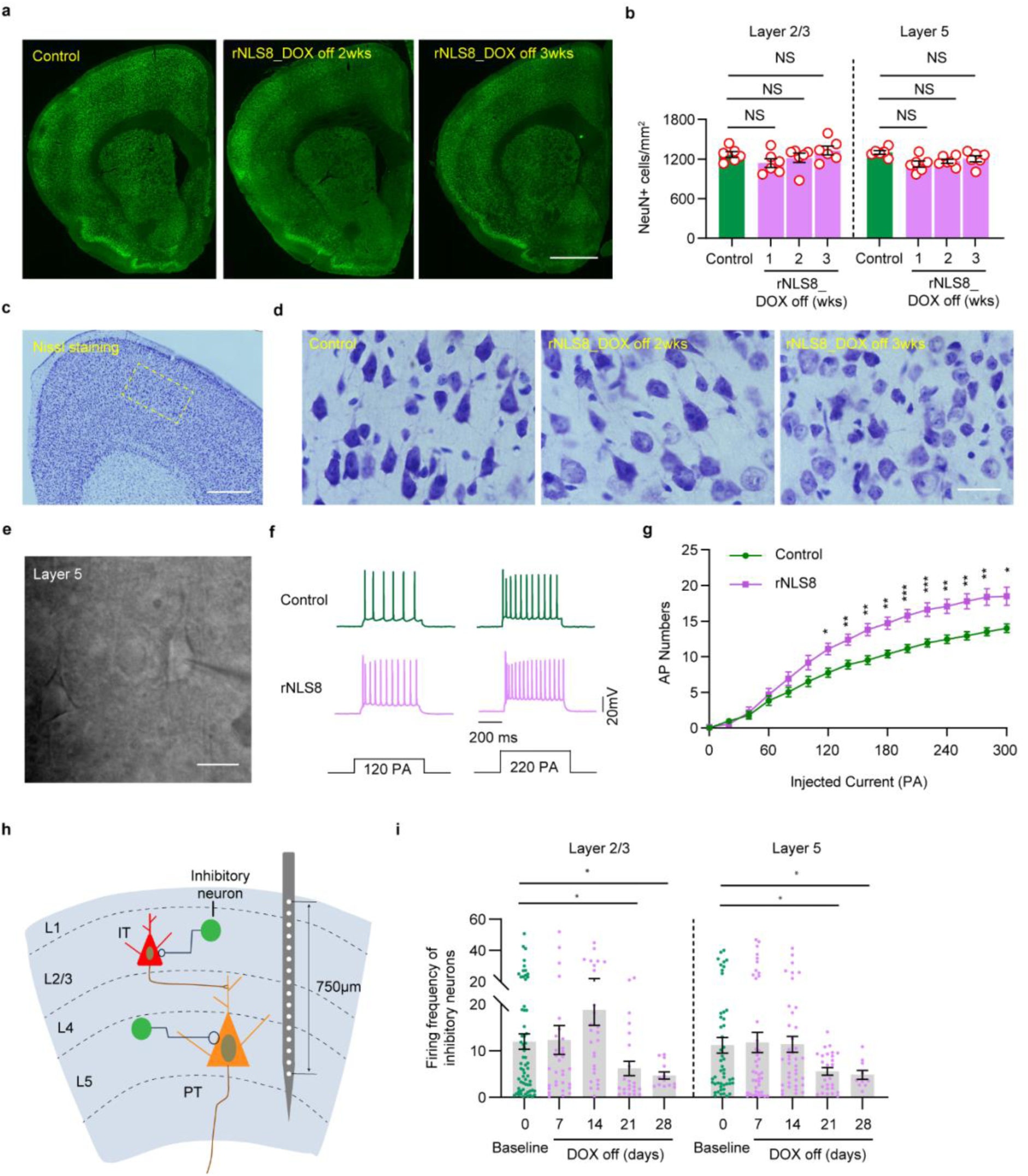
Cortical hyperactivity is associated with increased excitability of pyramidal neurons, related to Figure 1. **a,b,** Representative immunostaining images (**a**) and quantification (**b)** of NeuN (green) in M1 of control mice and rNLS8 mice at 2 weeks and 3 weeks post-DOX diet removal (*n* = 6 per group). Layer 2/3 and layer 5 are separatly quantified. Scale bar, 200 µm. **c,** Representative images of Nissl staining in M1 of a control mouse. A dotted white box indicates the motor cortical layer 5 area. Scale bar, 100 μm. **d,** Nissl staining of motor cortical layer 5 in control and rNLS8 mice at 2 weeks and 3 weeks post-DOX diet removal. Scale bar, 50 μm. **e,** Motor cortical layer 5 pyramidal neuron with a patch electrode for whole-cell recording. **f,** Electrophysiological profile of layer 5 pyramidal neurons in control and rNLS8 mice at day 16 post-DOX diet removal. The upper panel shows the action potentials elicited by the corresponding current step in the bottom panel. **g.** Quantification of action potentials in layer 5 pyramidal neurons elicited by corresponding current steps in control and rNLS8 mice at day 16 post-DOX diet removal (*n* = 23 neurons from 3 mice for the control group and *n* = 18 neurons from 3 mice for the rNLS8 group). **h,** Schematic illustration of in vivo recording of inhibitory neurons using a silicon probe with 32 channels in M1 of the rNLS8 mouse model. **i**, Firing frequency of inhibitory neurons in layers 2/3 (65 neurons) and 5 (50 neurons) in M1 of rNLS8 mice during baseline and disease progression (*n* = 7 mice). Statistical analysis, one-way ANOVA followed by Dunnett’s post hoc comparison to control (**b**), two-way ANOVA followed by Sidak’s post-hoc (**g**), or one-way ANOVA followed by Dunnett’s post hoc comparison to baseline (**i**). Error bars, mean ± s.e.m. NS = not significant, \**P* < 0.05; \*\**P* < 0.01; \*\*\**P* < 0.001; \*\*\*\**P* < 0.0001.

**Extended Data Fig 3.**
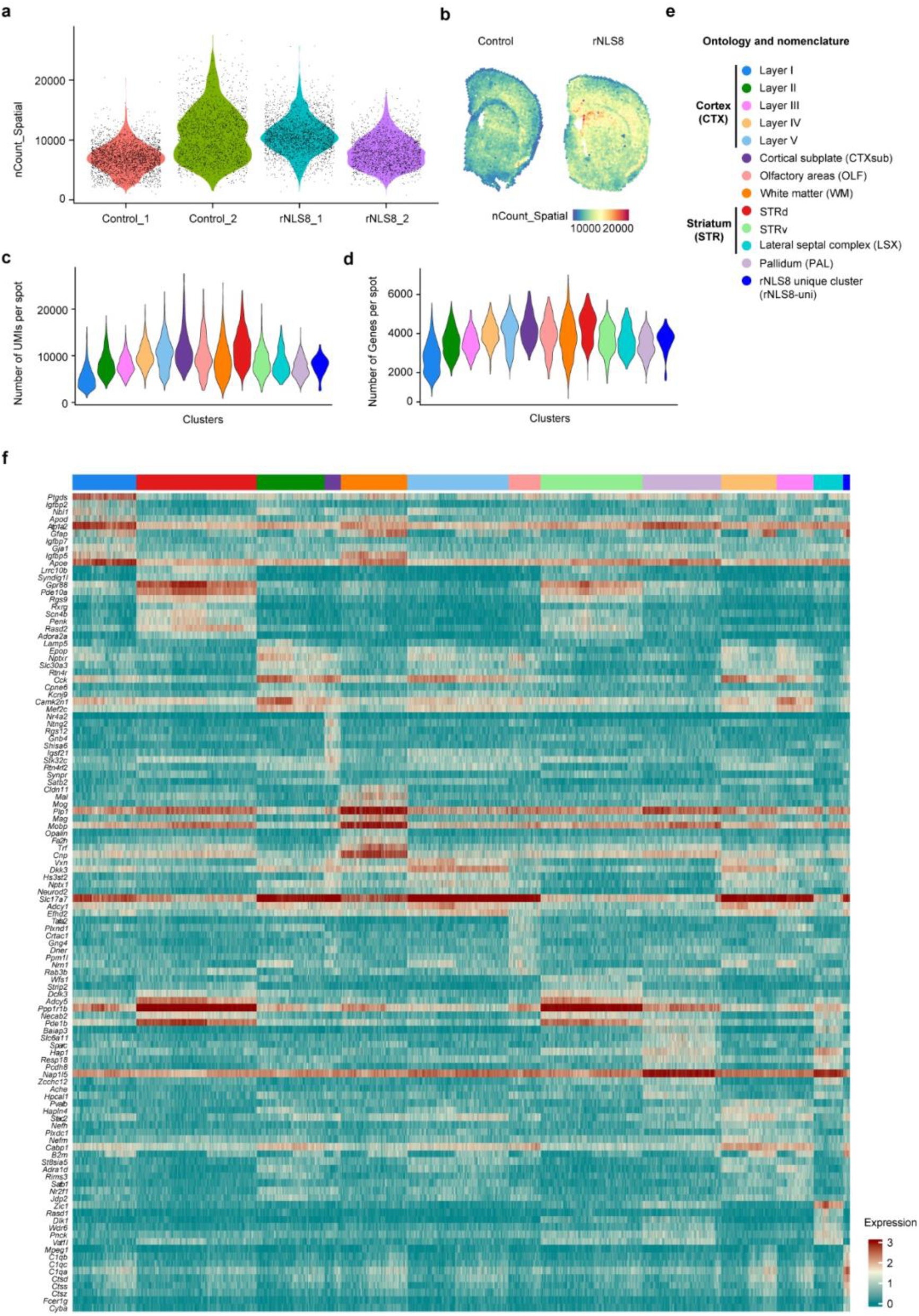
Basic characterization of spatial RNA sequencing, related to Figure 2. **a,** A violin plot showing the distribution of spots in the loaded data based on molecular counts per spot, allowing the exclusion of low-quality cells. **b,** The distribution of molecular counts across spots based on tissue anatomy. **c,d,** Violin plots of UMI count per spot (**c**) and genes per spot (**d**) in different regions. **e,** Complete data description and abbreviations of ontology and nomenclature for spatial transcriptome data. **f,** Heatmap of the top 10 genes in each cluster from spRNA-seq data.

**Extended Data Fig 4.**
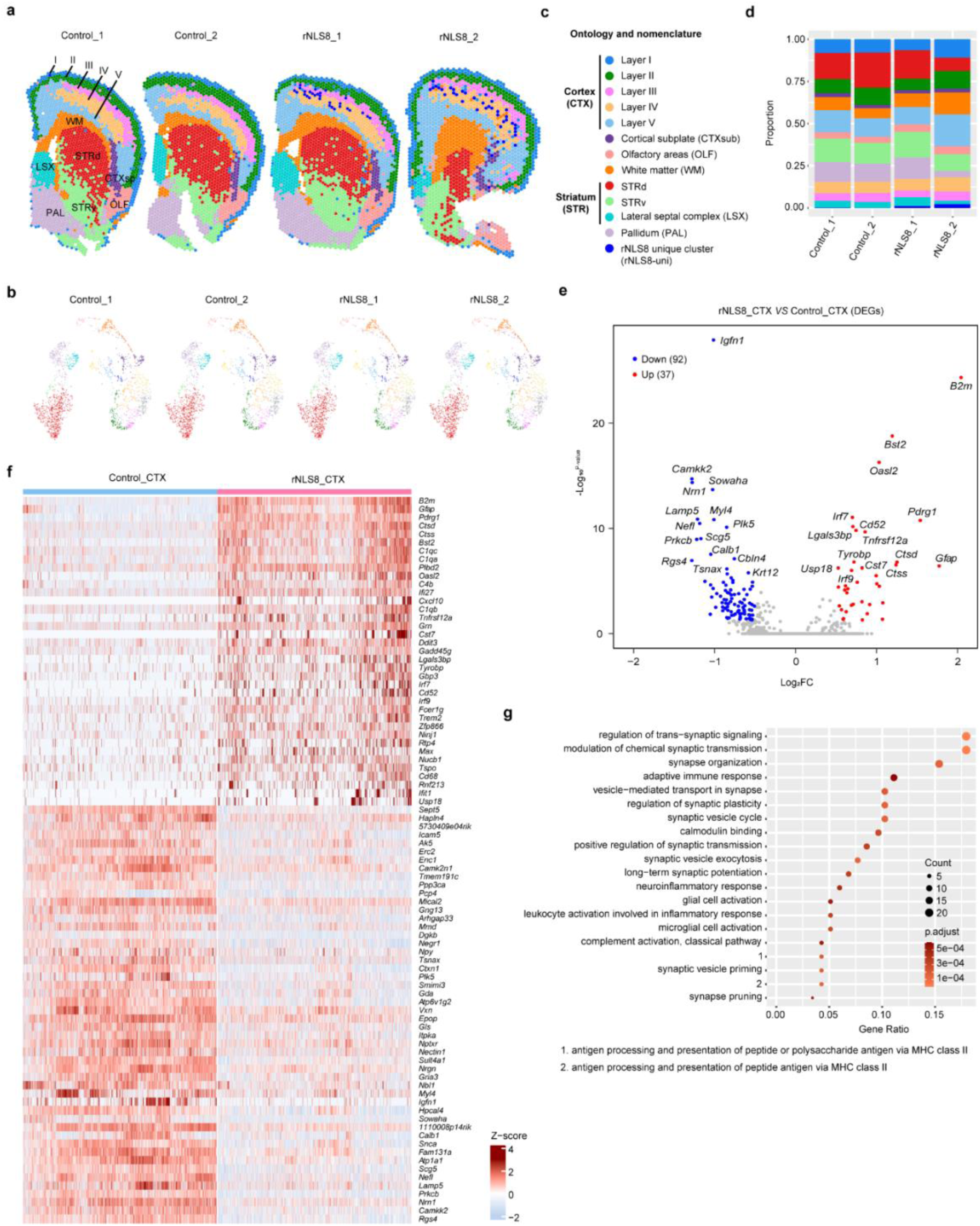
Spatial transcriptomics analysis of the cortex cluster, related to Figure 2. **a,b,** Spatial transcriptome data for individual subjects colored by cluster-level annotation and represented as spatial transcriptome (**a**) and UMAP (**b**). Cluster-level data were determined by Seurat clustering. **c,** Complete data description and abbreviations of ontology and nomenclature for spatial transcriptome data. **d,** Fraction of spots corresponding to each cluster. **e,** Volcano plot of DEGs in the ‘CTX cluster’ from spRNA-seq data. The CTX cluster (combining all cortical layers) in the control group served as a control. Significance is indicated at adjusted *P* value ≤ 0.05 and |avg_log2FC| ≥ 0.5 (log2FC, log2-fold change). **f,** Heatmap of DEGs in the ‘CTX cluster’ from spRNA-seq data. **g,** Gene Ontology (GO) enrichment analysis performed for the ‘CTX cluster’ from spRNA-seq data. Dot plot of the selected GO terms in order of gene ratio. The size of each dot indicates the number of genes in the significant DEG list that are associated with each GO term, and the color of the dots corresponds to the adjusted *P*-values.

**Extended Data Fig 5.**
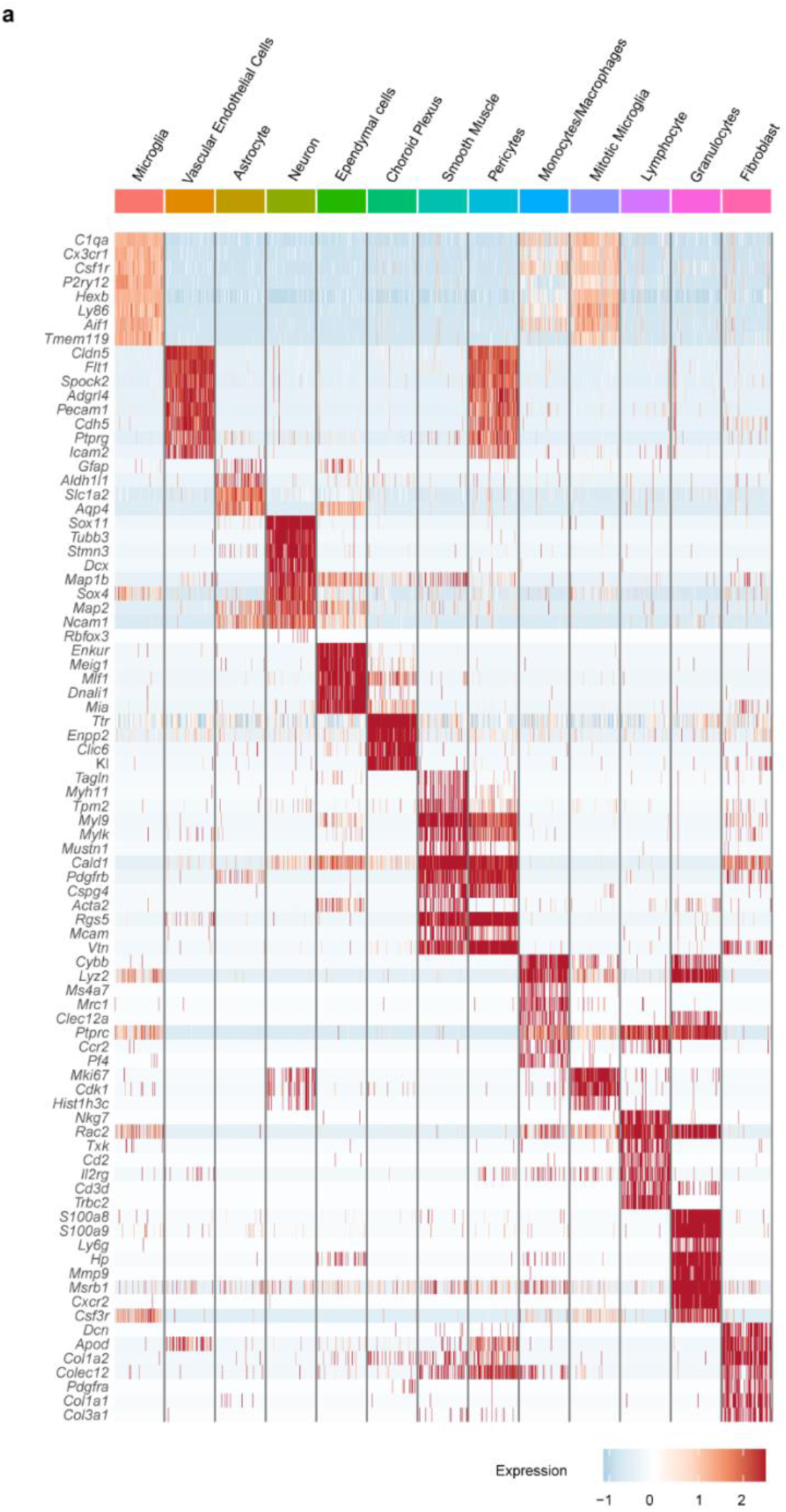
Heatmap of the marker genes in each cell type from single cell RNA sequencing, related to Figure 3. **a,** Heatmap of the marker genes in each cell type from scRNA-seq data.

**Extended Data Fig 6.**
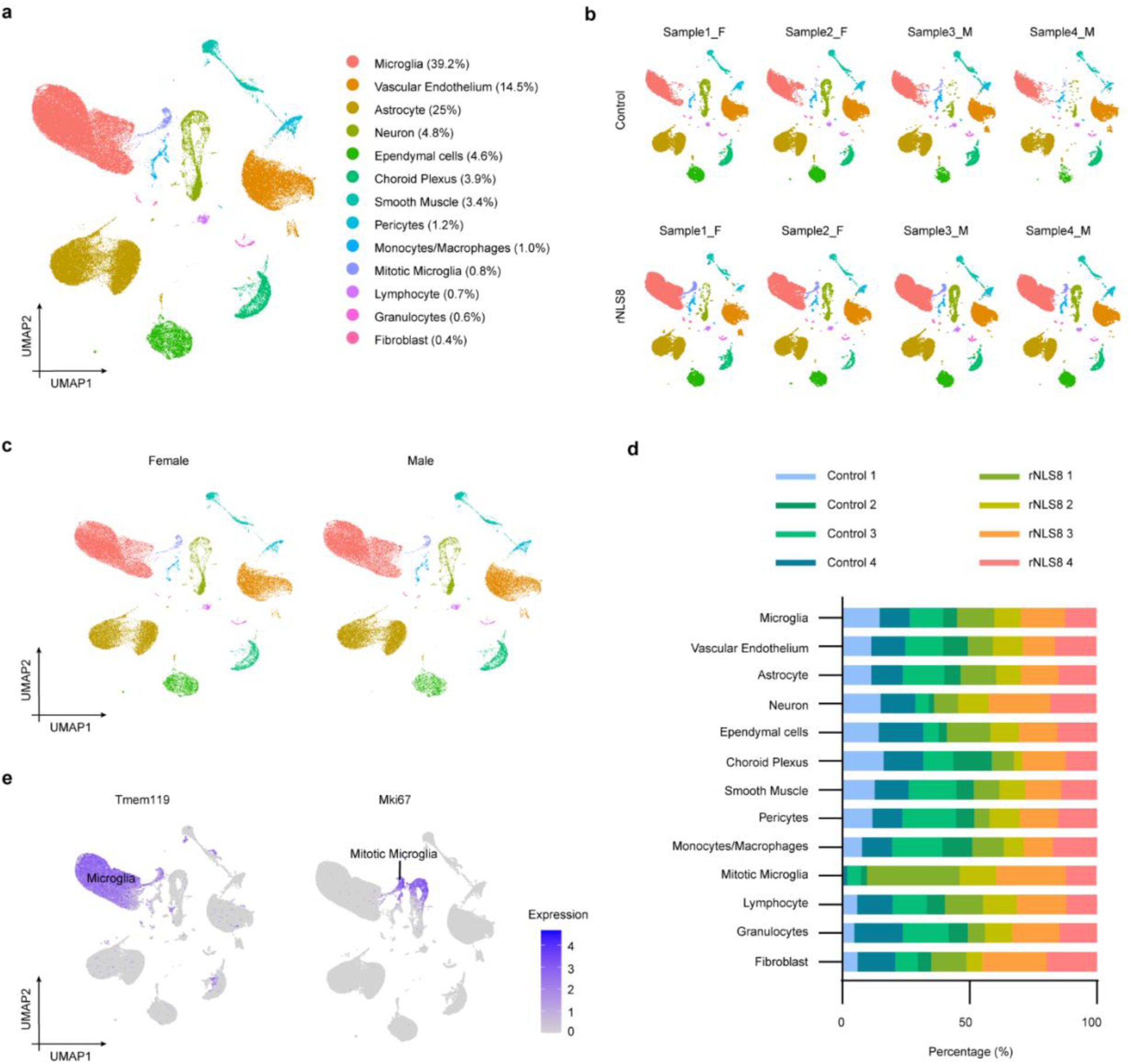
Basic characterization of single cell RNA sequencing, related to Figure 3. **a,** UMAP visualization of all cells identified by scRNA-seq. Cells are color-coded by their identities (number of cells = 101,162 from 8 samples). **b,** UMAP visualization of all cells identified by scRNA-seq in individual samples. **c,** UMAP visualization of all cells identified by scRNA-seq in female and male mice. **d,** Composition of individual samples within each cluster from scRNA-seq data. **e,** Feature plots of canonical markers defining microglia (*Tmem119*) and mitotic microglia (*Mki67*) identified from scRNA-seq analysis.

**Extended Data Fig 7.**
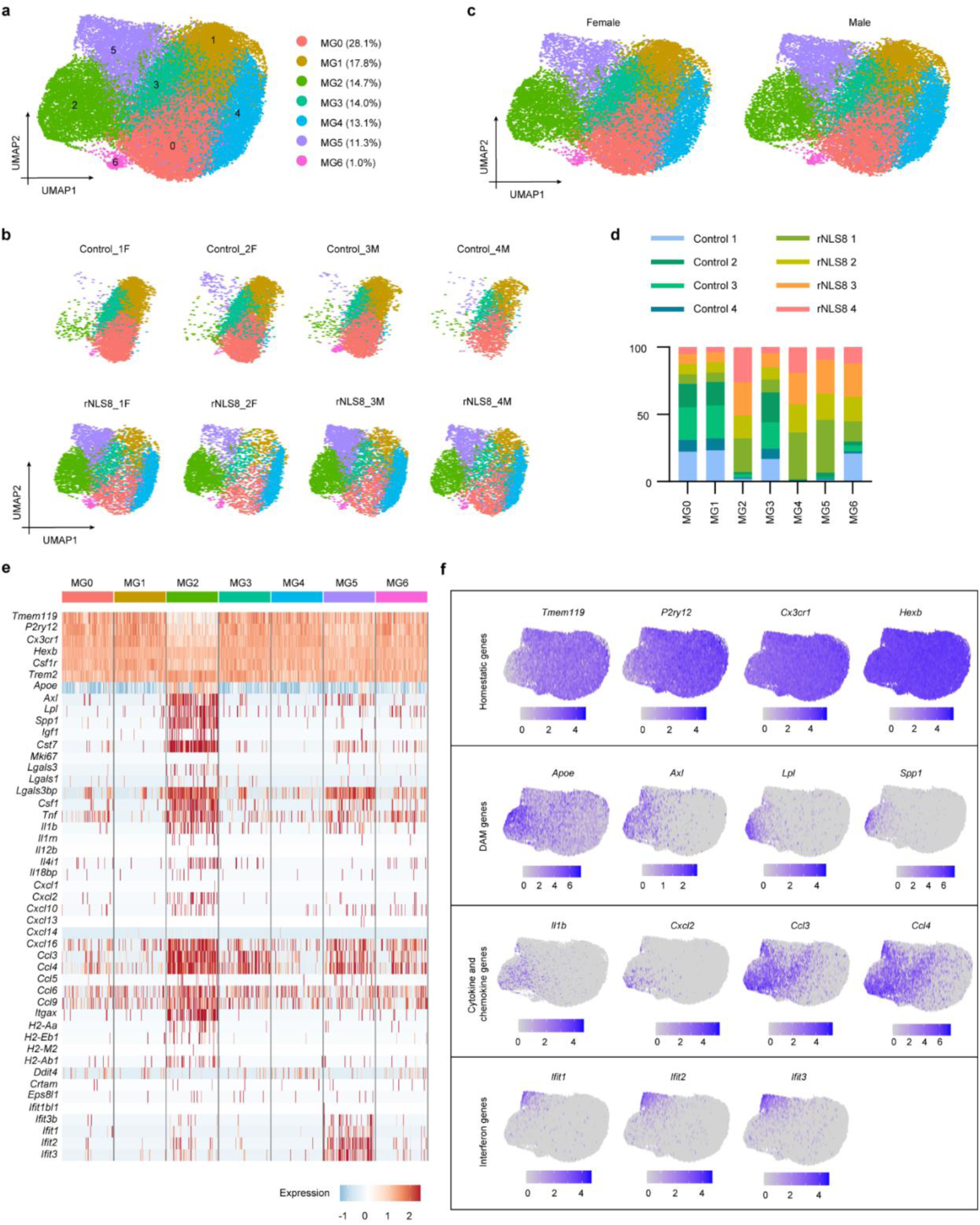
Basic characterization of microglial subclusters from single cell RNA sequencing analysis, related to Figure 4. **a,** UMAP visualization of microglia subclusters identified by scRNA-seq. Cells are color-coded by their identities (number of cells = 39,645 from 8 samples). **b,** UMAP visualization of microglia subclusters identified by scRNA-seq in individual samples. **c,** UMAP visualization of microglia subclusters identified by scRNA-seq in female and male mice. **d**, Individual sample composition of each microglia subcluster from scRNA-seq data. **e,** Heatmap of the marker genes in each microglia subcluster from scRNA-seq data. **f,** Feature plots showing the expression of selected markers for microglial subtypes.

**Extended Data Fig 8.**
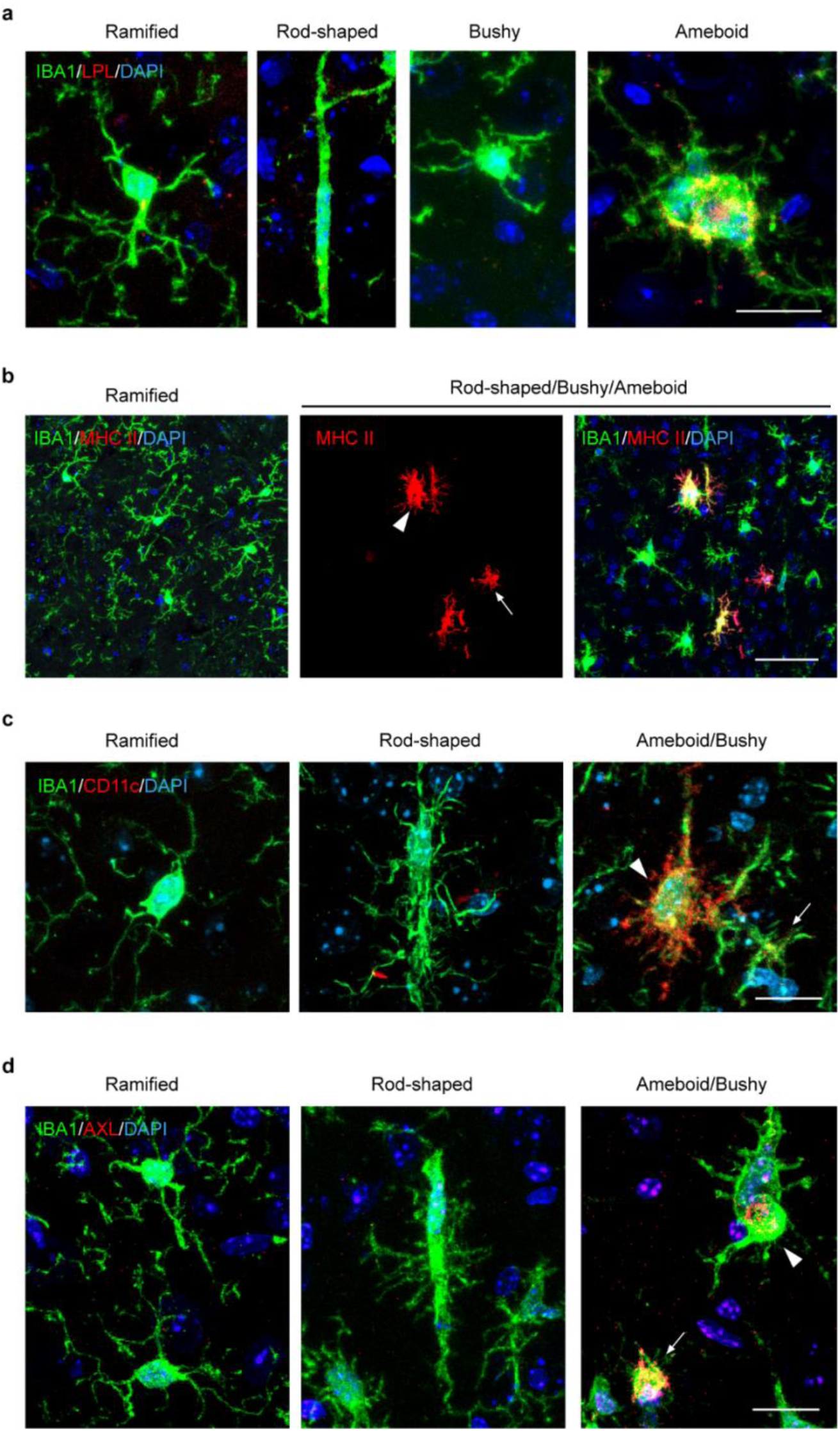
Marker screening for rod-shaped microglia, related to Figure 4. **a-d,** Representative images of LPL (**a**, red, Scale bar, 20 µm), MHC II (**b**, red, Scale bar, 50 µm), CD11c (**c**, red, Scale bar, 20 µm), and AXL (**d**, red, Scale bar, 20 µm) expression in ramified, rod-shaped, bushy, and amoeboid microglia (IBA1, green) at 3 weeks post-DOX diet removal. Bushy microglia are indicated by white arrows. Amoeboid microglia are indicated by white arrowheads.

**Extended Data Fig 9.**
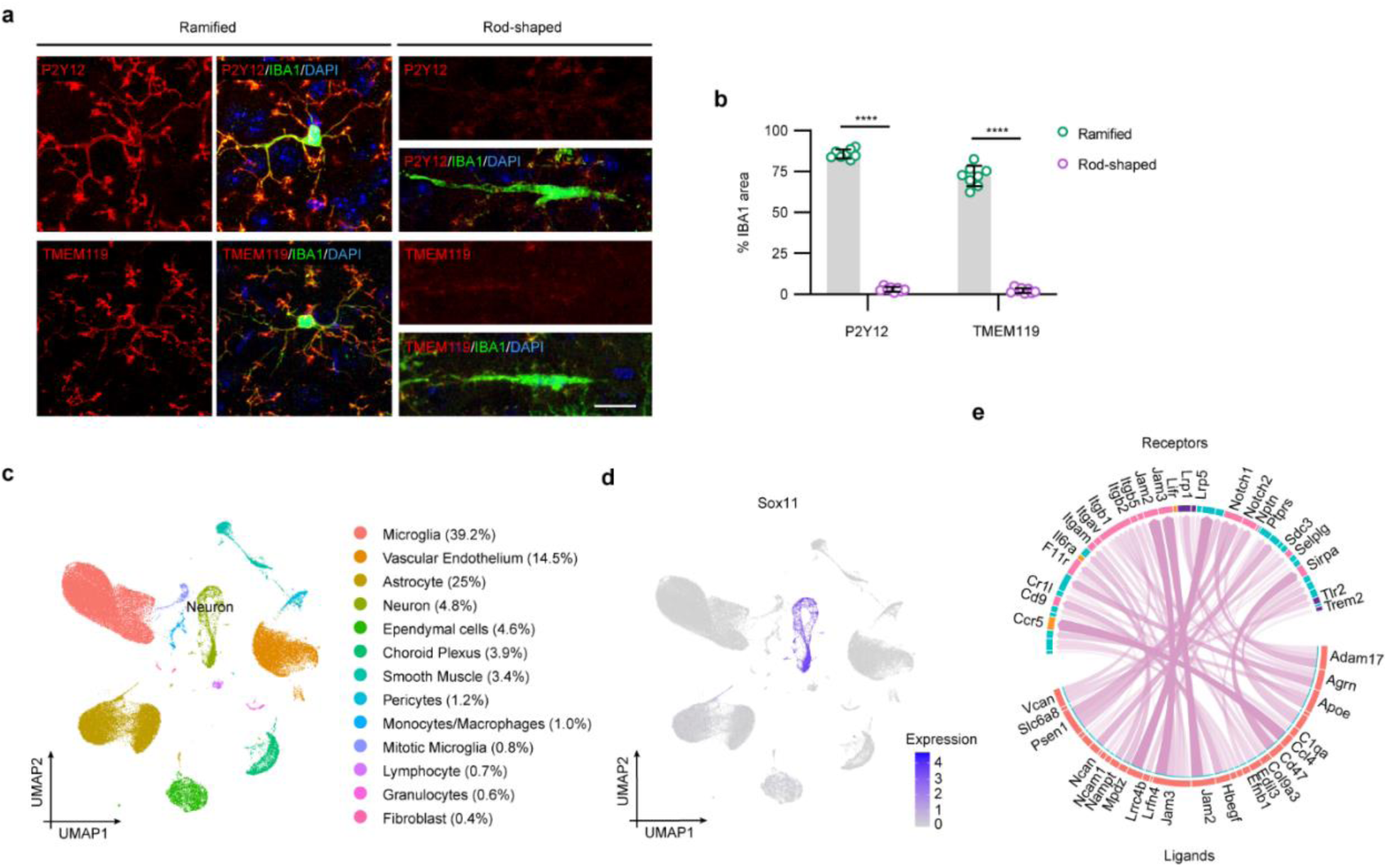
Characteristics of rod-shaped microglia and their interaction with neuronal dendrites, related to Figure 5. **a,** Representative images of P2Y12 and TMEM119 expression in ramified (control) and rod-shaped microglia (rNLS8) at 3 weeks post-DOX diet removal. Scale bar, 20 µm. **b,** Quantification of relative P2Y12 and TMEM119 positive areas in ramified (control) and rod-shaped microglia (rNLS8) at 3 weeks post-DOX diet removal. **c,** UMAP visualization of all cells identified by scRNA-seq. Cells are color-coded by their identities. **d,** Feature plots of canonical markers defining neurons (Sox11). **e,** Circos plot depicting links between ligands (in neuron) and their receptors (in microglia). The width and opacity of the links correlate with ligand-receptor interaction weights and ligand activity scores, respectively. Statistical analysis, two-tailed unpaired Student’s *t*-test (**b**). Error bars, mean ± s.e.m. NS = not significant, \**P* < 0.05; \*\**P* < 0.01; \*\*\**P* < 0.001; \*\*\*\**P* < 0.0001.

**Extended Data Fig 10.**
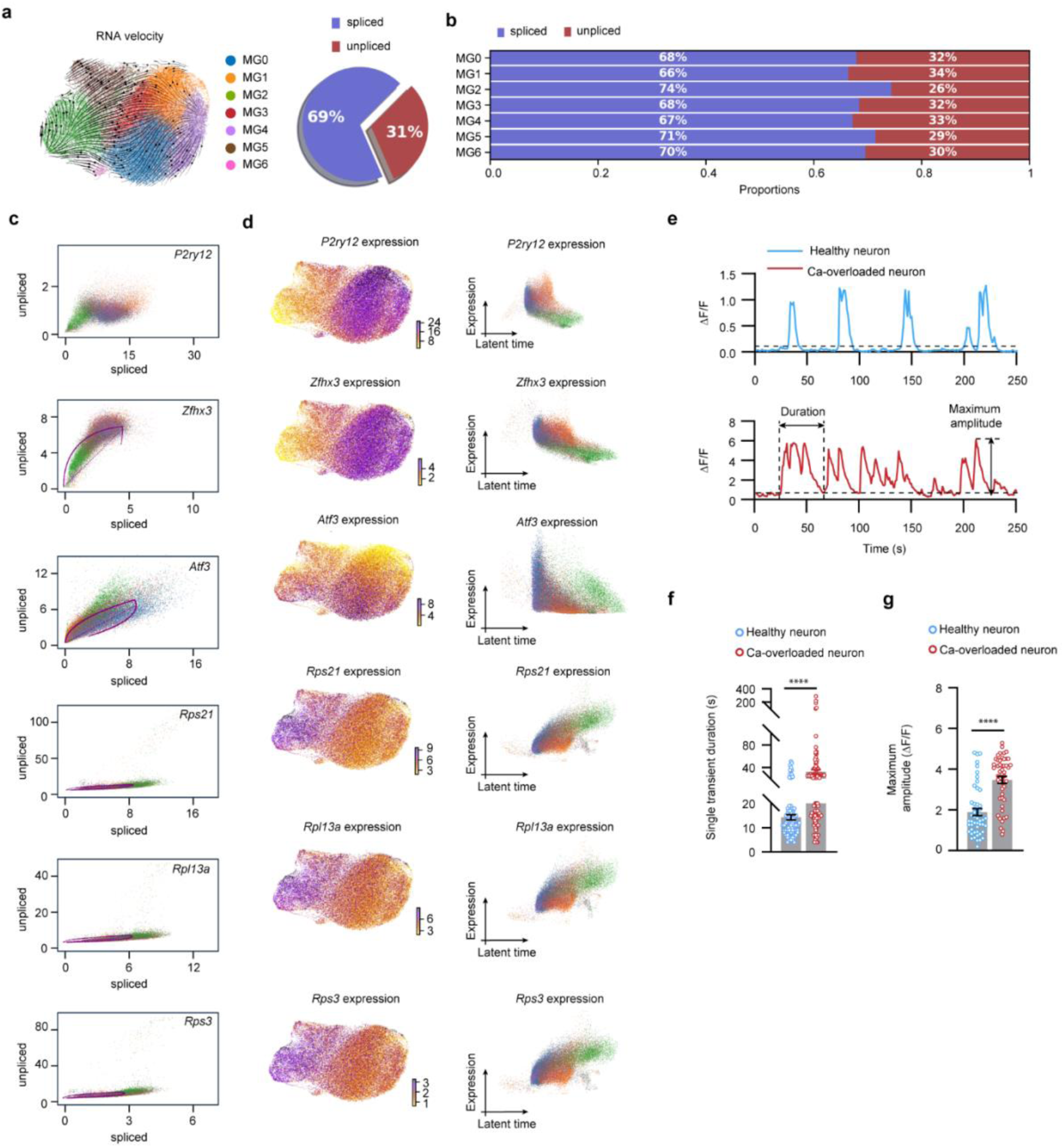
Basic characterization of RNA velocity analysis from single cell RNA sequencing data, related to Figure 6. **a,** Left: RNA velocity derived from the dynamical model for microglia subclusters visualized as streamlines in a UMAP-based embedding with scVelo. Right: Pie chart showing the proportions of spliced and unspliced (intron-containing) sequencing reads in the microglia cluster (combined from all samples). **b,** Bar chart showing the proportions of spliced and unspliced (intron-containing) sequencing reads in each of the microglia subclusters. **c,** Phase portraits of selected genes. **d,** The expression plots (left) and referred expression levels along latent time (right) of selected genes. **e,** ΔF/F calcium traces from the soma of a representative healthy neuron at baseline and neurons with Ca overload at 16 days post-DOX diet removal with the threshold line (black dotted line). Duration of single transient and the maximum amplitude are indicated by black arrows. **f,** Quantification of the duration of single transients in healthy neurons and neurons with Ca overload. **g,** Quantification of the maximum amplitude of Ca activity in healthy neurons and neurons with Ca overload. Statistical analysis, two-tailed unpaired Student’s t-test (**f and g**). Error bars, mean ± s.e.m. NS = not significant, \**P* < 0.05; \*\**P* < 0.01; *** *P* < 0.001; **** *P* < 0.0001.

